# Engram cell connectivity as a mechanism for information encoding and memory function

**DOI:** 10.1101/2023.09.21.558774

**Authors:** Clara Ortega-de San Luis, Maurizio Pezzoli, Esteban Urrieta, Tomás J. Ryan

**Affiliations:** School of Biochemistry and Immunology, Trinity College of Dublin, Dublin, Ireland; Trinity College Institute of Neuroscience, Trinity College Dublin, Dublin, Ireland; Brain Mind Institute, School of Life Sciences, École Polytechnique Fédérale de Lausanne, Switzerland; Florey Institute of Neuroscience and Mental Health, Melbourne Brain Centre, University of Melbourne, Melbourne, Victoria, Australia; Child & Brain Development Program, Canadian Institute for Advanced Research (CIFAR), Toronto, Ontario, Canada

## Abstract

Information derived from experiences is incorporated into the brain as changes to ensembles of cells, termed engram cells, that allow memory storage and recall. The mechanism by which those changes hold specific information is unclear. Here we test the hypothesis that the specific synaptic wiring between engram cells is the substrate of information storage. First, we monitor how learning modifies the connectivity pattern between engram cells at a monosynaptic connection involving the hippocampal vCA1 region and the amygdala. Then, we assess the functional significance of these connectivity changes by artificially activating or inhibiting its presynaptic and postsynaptic components respectively. Finally, we identify a synaptic plasticity mechanism mediated by PSD-95, which impacts the connectivity pattern among engram cells and contributes to the long-term stability of the memory. These findings impact our theory of learning and memory by helping us explain the translation of specific information into engram cells and how these connections shape brain function.

## INTRODUCTION

The brain’s memory systems encode information that allows animals to adapt their behaviour to a constantly changing environment ^1–3^. Learning is the process by which experiences are translated into lasting changes in the brain to enable behavioural adaptation. Engram cells are believed to be a subset of neurons that are activated by experiences, undergo molecular and structural modifications to encode a memory, and are necessary and sufficient to elicit a behavioural response when reactivated ^4^. The development and application of engram labelling technology has allowed the functional and molecular characterization of the engram cells that contribute to the storage of specific long-term memories ^5–7^. Artificial optogenetic activation of engram cells elicits the reactivation of a memory ^5,8^ and allows its manipulation ^8–11^, whereas its inhibition prevents the recall of the memory ^12,13^.

Though the behavioural functionality of engram cells has been well characterised, the molecular and cellular mechanisms that enable the formation of engram cell ensembles are largely unknown ^14–16^. Plasticity mechanisms can modify both *synaptic weights*, the strength of the connection between cells, and *synaptic wiring*, the structural patterns of connections that are formed between cells ^17^. Both mechanisms are likely involved in engram cell formation and function ^18^. Engram cells exhibit higher AMPA to NMDA ratio, more dendritic spines, and an enhanced number of synaptic inputs ^10,19–22^. Structurally, connectivity patterns between engram cells distributed across different brain areas are modified by learning ^23^. The analysis of the downstream effect of engram cell activation serves as a readout for engram cell connectivity, and this mechanism has been implicated as a substrate of memory persistence ^10,24,25^. It can be hypothesised that these stable connectivity patterns between engram cells allow engrams to survive amnesia ^8,10,26–31^ and are behind the creation of false or artificial memories ^8,32^.

Projections from the ventral CA1 (vCA1) subregion in the hippocampus to the basal amygdala (BA) are implicated in contextual fear memories ^33^, and selective strengthening of synaptic connections between vCA1 to BA neurons has been associated with memory encoding ^22^. We capitalised on this monosynaptic connection to investigate how changes of synaptic wiring that modify the engram connectivity pattern can explain information specificity in the process of learning and encoding.

Several molecular mechanisms adapt synaptic plasticity in response to neuronal activation. Postsynaptic density protein 95 (PSD-95) is a postsynaptic scaffolding protein that determines synaptic function, including synaptic strength and maturation ^34–37^. Dysregulation of PSD-95 in knockout models increases long-term potentiation and provokes a deficit in spatial learning ^35,37^. However, engram specific molecular manipulations are needed to understand how these synaptic plasticity mechanisms translate into information specificity.

Here we test the hypothesis that information encoding and memory function depends on the specific connectivity pattern formed between engram cells. First, we monitor how the engram synaptic wiring changes in response to an experience in a monosynaptic preparation between vCA1 and BA. Then we evaluate the functional role of the change in the diagram of connections by artificial activation and inhibition of its components and finally, we identify a synaptic plasticity mechanism mediated by PSD-95 that influences the connectivity pattern between engram cells and the long-term persistence of the memory.

## RESULTS

### Learning modifies synaptic wiring of engram cells

To study the effect of experience on the connectivity pattern between engram cells, we designed a behavioural paradigm that allowed us to monitor learning by following the updating of a neutral context memory after it is updated with aversive information (Figure 1, A). By re-exposure to a previously encoded and neutral context (Context A) and the immediate delivery of an electric shock (immediate shock), only animals with a pre-formed engram for A updated such engram with the new aversive information, and context A became A’. 24 hours after the updating, animals showed context-specific freezing behaviour to A’, indicating the formation of a fearful memory, specific to context A’ (Figure 1, B). The same aversive experience was unable to elicit freezing in the control group that lacked a pre-existing engram for the context (Not Linked I control group). Similarly, re-exposure to the context alone did not induce a fear response (Not Linked II control group). All animals were able to encode and recall a fearful memory (Fig 1, B, C) while fear generalisation did not extend to a neutral context (Context C, Fig 1, B). Finally, a 30 minute exposure to context A’ in absence of footshocks induced extinction of the fear memory, evidenced by a significant decrease in freezing behaviour (Suppl Fig 1).

**Figure 1.**
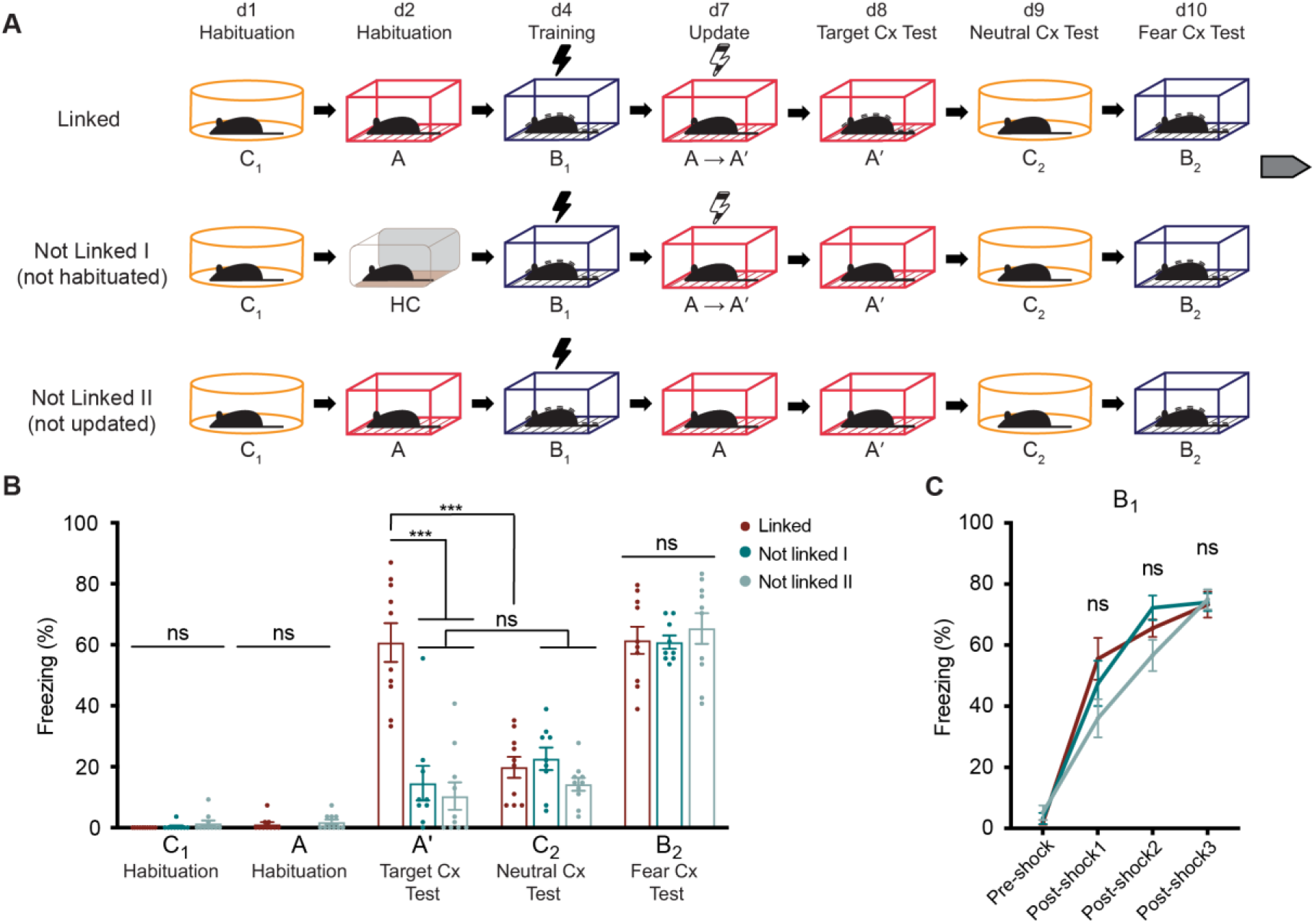
Animals learn to update a neutral context by forming a fearful memory while maintaining context discrimination. **A.** Behavioural schedule along the span of 10 days involving three separate contexts (A, B, C). Experimental group (Linked) was habituated to Context C and Context A, and then trained in Context B (contextual fear conditioning paradigm, black lightning symbol). Training paradigm involved 3 min exploration of context and the 1 min-spaced delivery of 3 foot-shocks. On day 7, during the Update of Context A, animals returned to A and immediately received a single foot-shock (immediate-shock paradigm, black and white lightning symbol), therefore updating their original engram for A and forming a fear association to A (A became A’). Presence of a fearful memory for the three contexts was tested on day 8 (A’), day 9 (C_2_) and day 10 (B_2_). Control group Not Linked I animals lack an existing engram for A to update, since they were not exposed to A prior to behavioural update and remained in their homecage (HC). Control group Not Linked II animals were not updated to A’ as foot-shock was not delivered during the corresponding session. **B.** Freezing behaviour as a readout for memory recall along the behavioural schedule. High freezing in A’ in Linked group but not in Not Linked I (not habituated) control indicates that animals updated their pre-existing engram for A to encode the new fearful information of A’, since Not Linked I group control, which had never been exposed to A were unable to form such fearful association. All groups showed context discrimination evidenced by low and high levels of freezing to Context C and B respectively during recall (C_2_ and B_2_). **C.** Training curve in Context B showing acquisition of a fearful memory evidenced by a freezing response during contextual fear conditioning in the three groups. Data presented as mean ± SEM. n = 9 - 10. Repeated measures two-way ANOVA, Tukey; *** p<0.001. ns, not significant. Cx, context; HC, homecage; Grey arrow indicates continuation in Suppl Fig 1.

To identify a change in synaptic wiring of engram cells for context A when it becomes A’, we hypothesised that the aversive experience would update the engram by recruiting new cells, and after doing so, these cells would partially overlap with the engram cells for a separate aversive experience of a similar nature (context B) encoded 3 days before. Therefore, to characterise the changes in synaptic wiring by learning, we used engram labelling technology to tag the BA engram component of experience B by local delivery of AAV_9_-c-fos-tTA and AAV_9_-TRE-RFP vectors (Fig 2, A - C). After updating A to A’, recall of A’ induced reactivation of engram B cells (Fig 2, D - F), measured by c-Fos expression, indicating a synaptic wiring change and a functional link (Figure 2). As a control, we showed that without the updating paradigm, these two contexts are represented as separate engrams in the BA (Suppl Fig 2).

**Figure 2.**
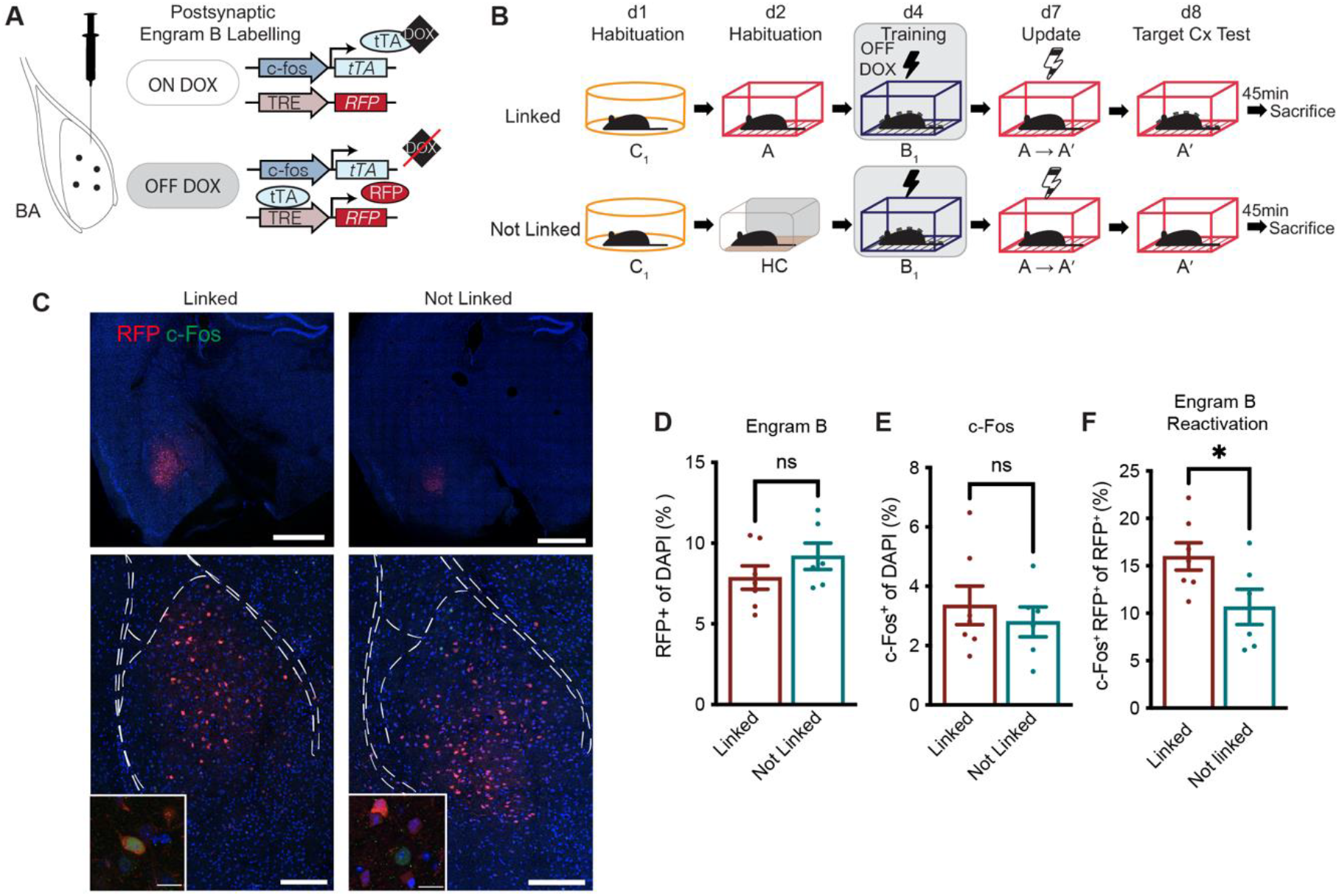
Learning modifies synaptic wiring of engram A cells after updating. A. Doxycycline-dependent engram labelling strategy to label engram B in the BA through viral stereotactic injection. **B.** After habituation to C and A on days 1 and 2 respectively, doxycycline was removed from the diet and engram B was labelled during training on day 4 (contextual fear conditioning paradigm, black lightning symbol). On day 7, engram A was updated to A’ (immediate shock paradigm, black and white lightning symbol). On day 8, engram B reactivation induced by A’ context was measured by sacrificing mice 45 min after a natural recall of A’. **C.** Representative images of red fluorescent protein (RFP, red), c-Fos (green) and overlap of the two signals in the BA. DAPI signal is represented in blue. Scale bars: 1 mm (upper panels), 200 µm (lower panels), insets: 20 µm. **D.** Engram B size in BA (RFP^+^ cell counts). **E.** c-Fos^+^ cell counts in BA. **F.** Engram B reactivation induced by natural recall of A’. RFP^+^ and c-Fos^+^ cell counts, measured as ratio of double positive cells per total of RFP^+^ cells. Data presented as mean ± SEM. n = 6 - 7. Unpaired t-test; * p<0.05. ns, not significant; BA, basal amygdala; Cx, context, DOX, doxycycline.

### Optogenetic stimulation of vCA1 neutral engram cells elicits updated memory recall and downstream reactivation of engram B cells

In order to better understand how the connectivity pattern of engram A changes after the updating of the memory, we next focused on the monosynaptic connection between vCA1 and BA. First, we demonstrated the presence of a monosynaptic connection by detecting evoked postsynaptic currents (eEPSCs) in the BA. Optogenetic activation of vCA1, of both engram cells and non-engram cells specifically, induced a response in BA neurons (Suppl Fig 3).

To further confirm a change in the synaptic wiring in engram A, we tagged the presynaptic element of the connection (vCA1) that encodes the contextual component of the experience, during the encoding of the neutral context. To do so, we used an engram-specific labelling system based on the tamoxifen-inducible method where we locally injected AAV_9_-DIO-ChR2- EGFP vector into TRAP2 animals ^38,39^ (Fig 3, A, C). We combined this strategy with the DOX-dependent system in the BA to tag the engram B cells (Fig 3, F, G). After the update of context A, we artificially activated the presynaptic component of engram A in a neutral context (Fig 3, B). Artificial activation of vCA1 engram A cells was sufficient to elicit a behavioural response in the animals (Fig 3, D - E) as well as a downstream activation of engram B (Fig 3, H - J). Reactivation of B induced by artificial activation of A in experimental animals, but not control animals, indicates a change in synaptic wiring upon learning. Taken together, these results demonstrate that the connectivity pattern between engram cells has been modified in response to experience.

**Figure 3.**
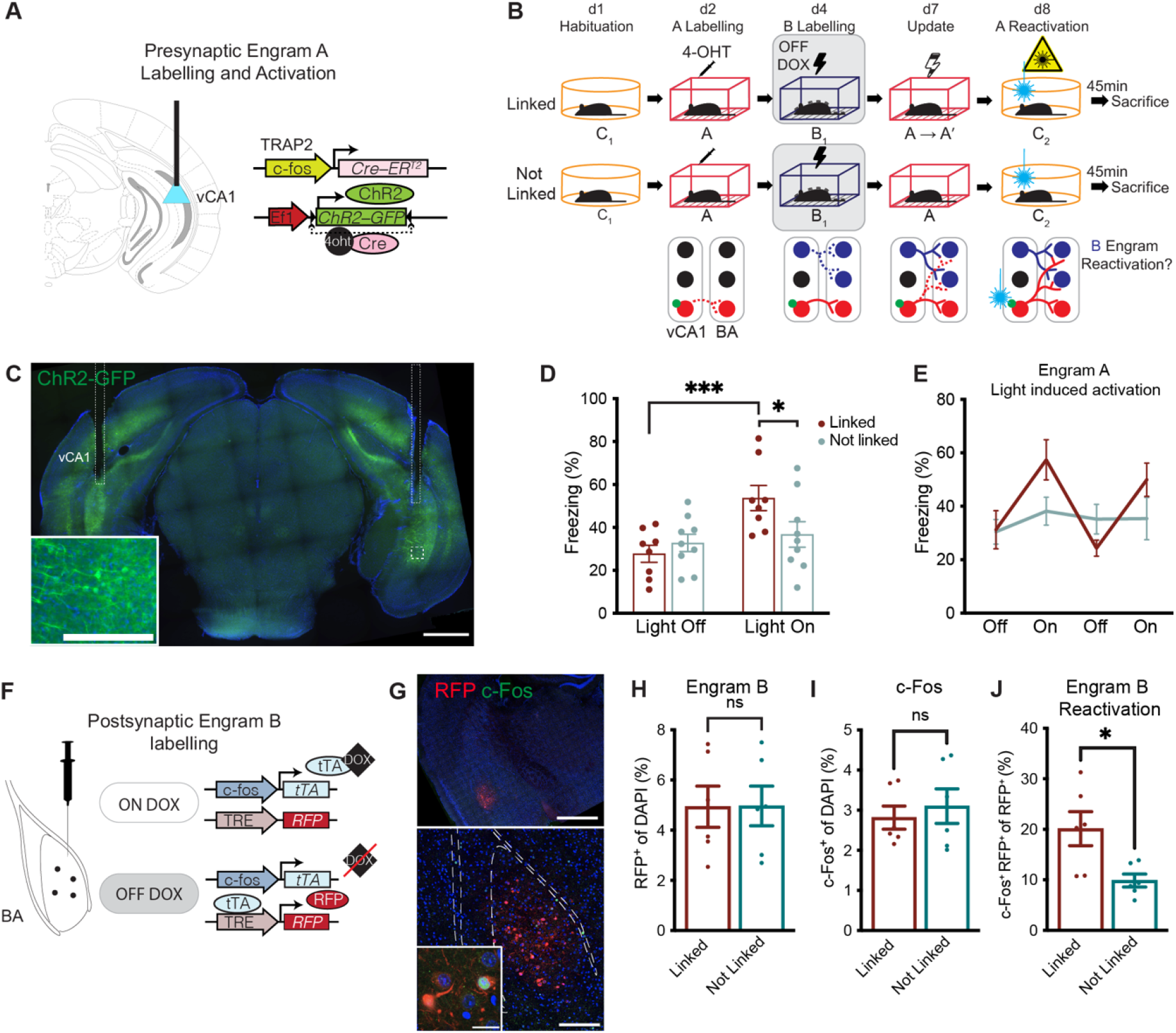
Optogenetic stimulation of vCA1 neutral engram cells elicits updated memory recall and downstream reactivation of engram B cells A. Tamoxifen-induced engram labelling strategy to allow optogenetic activation and tagging of engram A by AAV-delivery of a Cre-dependent chanelrhodopsin (ChR2) protein and EYFP reporter into the vCA1 of TRAP2 mice. Optic fibers were implanted to allow the delivery of light. **B.** Upper panels: behavioural schedule. Bottom panels: diagrams representing the connection vCA1 to BA. Animals were habituated to context C on day 1. On day 2, labelling of engram A was achieved by the injection of tamoxifen (4- OHT) immediately after habituation to A. A neutral engram for context A was formed after the experience (red cells in the diagrams, dotted line represents new connections) and its presynaptic component was labelled with ChR2 (green circle in the diagrams). Doxycycline (DOX) was removed from the diet of the animal and on day 4 engram B was labelled during training (contextual fear conditioning paradigm, black lightning symbol) in BA. An engram for a fearful experience for B was formed and its postsynaptic component was labelled with red fluorescent protein (RFP) in the BA (blue cells in the diagrams, dotted lines represent new connections). On day 7, engram A is updated to A’ (immediate shock paradigm, black and white lightning symbol). This experience modifies the wiring of engram A, creating new connections between the presynaptic vCA1 component of original neutral engram A and the postsynaptic BA component of fearful engram B. On day 8, presynaptic vCA1 component of engram A was artificially activated by delivery of light (blue laser beam symbol, yellow laser symbol), while mice explored neutral context C. Finally, engram B reactivation induced by artificial activation of A’ was measured by sacrificing mice 45 min after optogenetic stimulation. **C.** Representative images of EYFP signal (green) in vCA1 and implant location. DAPI signal is represented in blue. Scale bars: 1000 µm, inset: 250 µm. **D-E.** Light-induced activation of A’ in vCA1 elicited freezing while in Context C in experimental (Linked) but not control (Not linked) groups during light-on periods. **F**. DOX-dependent engram labelling strategy used to label engram B in the BA in the same mice. **G**. Representative images of red fluorescent protein (RFP, red), c-Fos (green) and overlap of the two signals in the BA. DAPI signal is represented in blue. Scale bars: 1mm (upper panel), 200 µm (middle panel), inset: 25 µm. **H.** Engram B size in BA (RFP^+^ cell counts). **I.** c-Fos^+^ cell counts in BA. **J.** Engram B reactivation induced by optogenetic recall of A’. RFP^+^ and c-Fos^+^ cell counts, measured as the ratio of double positive cells per total to RFP^+^ cells. Data presented as mean ± SEM. n = 6 - 9. Repeated Measures two-way ANOVA, Sidak multiple comparisons (D) or unpaired t-test (H - J); *p<0.05; ***p<0.001; ns, not significant. BA, basal amygdala.

### Synaptic wiring changes are required to update the memory

In order to address the functional relevance of the change in the connectivity pattern between engram cells for the updated memory expression, we tagged engram B cells in the BA with an inhibitory opsin, ArchT (Fig 4, A, C). After updating the memory, we asked whether the activity of the newly recruited cells in the BA, (engram B cells) was required for the recall of A’ (Fig 4, B). A significant decrease in freezing behaviour in A’ upon light-delivery, when compared to the No Light group was found, demonstrating that, at least partially, the recall of the updated memory requires the activation of the engram B cells (Fig 4, D). Importantly, optogenetic silencing of a non-meaningful engram in the BA of similar size (Fig 4, F) did not induce similar effects (Random ensemble group). As a control, all groups exhibited normal context discrimination, as well as normal recall of a fearful memory (Suppl Fig 4). This demonstrates that changes in synaptic wiring between engram cells are necessary for the updating of the memory.

**Figure 4.**
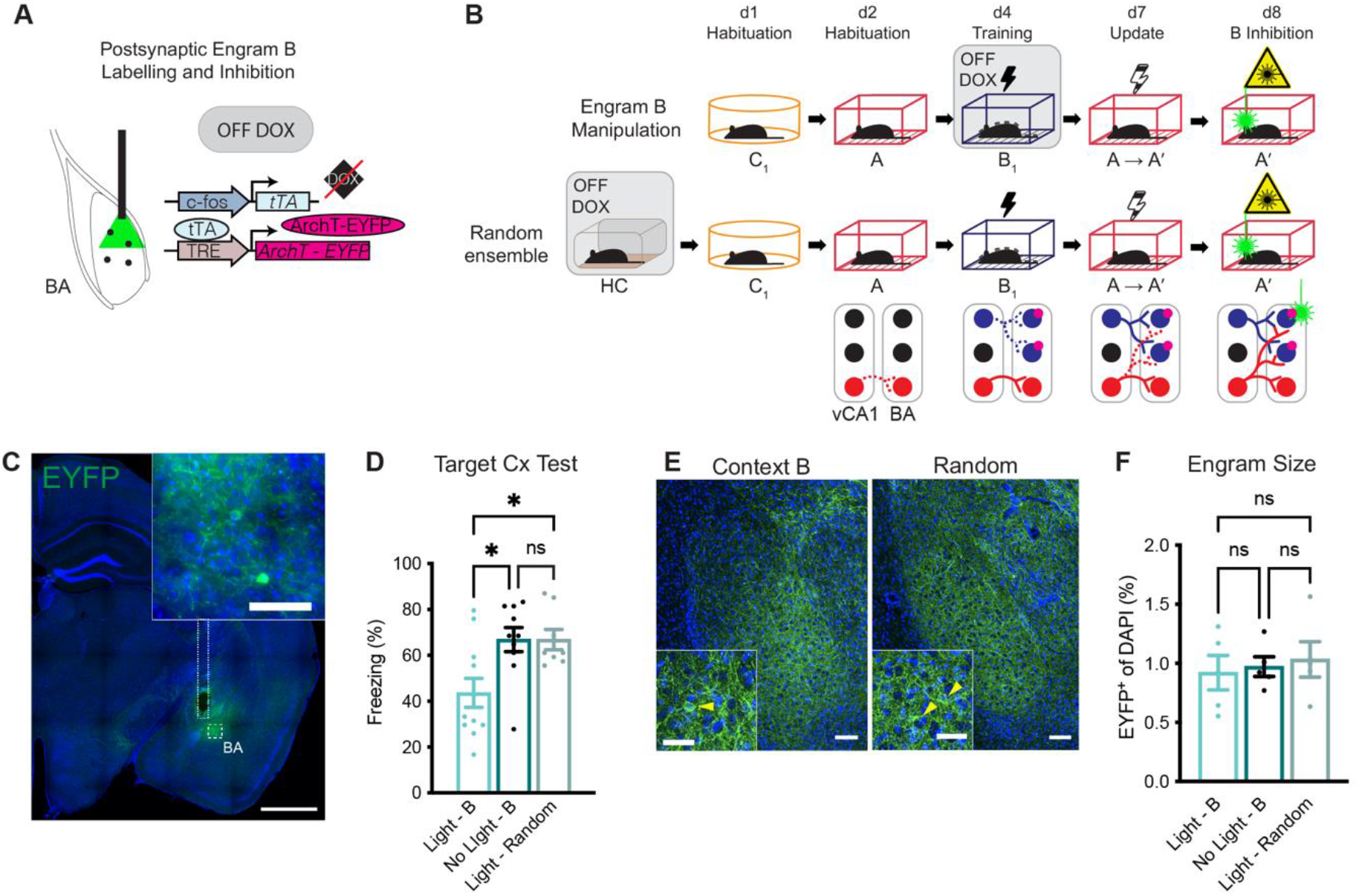
Synaptic wiring changes are required to update the memory. A. Doxycycline-dependent engram labelling strategy to allow optogenetic inhibition and tagging of engram B by AAV-delivery of a doxycycline (DOX) dependent archaerhodopsin (ArchT), an inhibitory opsin and fluorescent protein EYFP into the BA. Optic fibres were implanted to allow the delivery of light. **B.** Behavioural schedule. After habituation to context C and A on days 1 and 2 respectively, doxycycline was removed from the diet and engram B was targeted during training on day 4 (contextual fear conditioning paradigm, black lightning symbol) with ArchT (pink circle in the bellow diagram). To label a random ensemble, DOX was removed from the diet for 50 h (instead of the 36h period of the routine protocol, including two cage changes, see materials and methods for more details). On day 7, engram A was updated to A’ (immediate shock paradigm, black, lightning symbol). On day 8, activity of engram B was prevented by the delivery of light to the BA during recall of A’ (green laser beam symbol, yellow laser symbol). **C.** Representative images of EYFP signal (green) in BA and implant location. DAPI signal is represented in blue. Scale bars: 1000 µm, inset: 100 µm. **D.** Inhibition of engram B through light during recall of A’ reduced freezing levels (Light - B group) when compared to animals that received no light over engram B (No-light control, No Light -B), or to animals that received light over a random ensemble in the BA (Random ensemble, Light - Random). **E.** Representative images of EYFP signal (green) in BA of mice with engram B labelled (Context B, left) and mice with a Random ensemble labelled (Random, right). Scale bars: 100 µm, insets: 50 µm. Yellow arrowheads point at EYFP^+^ neurons. **F.** Size of engram B labelled in Light - B and No Light - B groups is similar to size of random ensemble (Light - Random), measured by EYFP^+^ cell counts in BA. Data presented as mean ± SEM. n = 8 - 11 (D), 5 (F). One-way ANOVA, Tukey multiple comparisons; *p<0.05; ns, not significant. BA, basal amygdala; Cx, context.

### Plasticity mechanisms associated with changes in synaptic wiring in engram cells

Next, to characterise how these changes in synaptic wiring correlate with parameters of synaptic plasticity, we analysed dendritic spines and synaptic markers. To assess a modification in dendritic spine densities and spine morphology, we analysed a battery of parameters related to dendritic spine formation in engram B cells (Fig 5, B - E; Suppl Fig 5, A - B). After recruiting afferent connections from the updated A’ engram, engram B cells showed an activity-dependent increase in the number of dendritic spines. However, 45 minutes after recall on day 7, the total number of synapses onto engram B cells, quantified by the number of events where proximal presence of the excitatory marker type I vesicular glutamate transporter 1 (VGLUT1) puncta to intra-engram postsynaptic density protein 95 (PSD-95) puncta were detected, did not change after activity (Fig 5, F - I). Although the number of PSD- 95^+^ puncta inside engram cells did not change, we found that the total expression of PSD-95^+^ decreased in recall-activated engram B neurons that received the newly recruited connections, 45 minutes after recall (Fig 5, J). Since synapses rely on a variety of postsynaptic scaffolding proteins for stabilization ^40,41^, we extended our analysis to Synapse-Associated Protein 102 (SAP102), a post-synaptic scaffolding protein that is present in a separate subset of synapses throughout the brain ^42^. Total number of VGLUT1^+^-SAP102^+^ synapses did not change after reactivation of engram B in the linked group (Suppl Fig 5, C - G), neither did the expression of SAP102. Finally, we explored whether the increase in engram B activation induced by recall of A’ in the updated group could be explained by a change in the inhibitory tone that these cells receive. The number of the postsynaptic inhibitory marker gephyrin (GPHN) puncta was decreased in engram B after recall of A’ (Fig 5, K - O). At the same time, the number of the presynaptic inhibitory marker vesicular GABA transporter (VGAT) puncta that engram B received, the number of inhibitory synaptic contacts measured by proximity of these two markers, or the total amount of Gephryn expression was not found to be altered.

**Figure 5.**
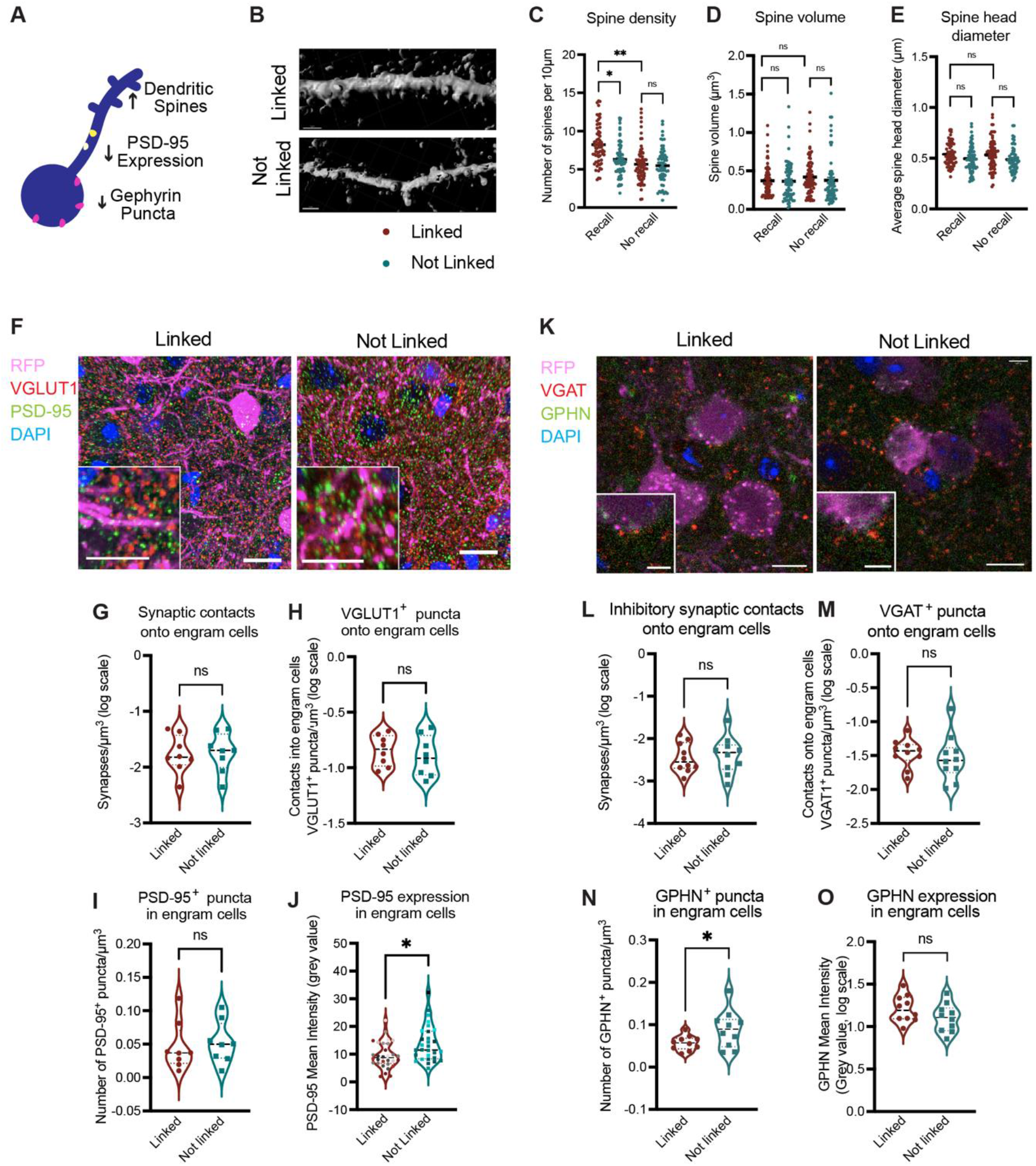
Plasticity mechanisms associated with changes in synaptic wiring in engram cells A. Diagram representing plasticity mechanisms associated with the reactivation of engram cells after synaptic wiring changes. **B-E.** Dendritic spine analysis of engram B in animals in the linked group (red) and the not linked group (green, not habituated control) after recall of A’ (Recall) or no recall (No recall). **B**. Representative image of a RFP^+^ dendrite as identified as a surface by the IMARIS software in each group. Scale bars: 3 µm. **C.** Spine density measured by number of spines in engram B cells. Average dendritic spine volume (**D**) and spine head diameter (**E**) per each 10 µm dendritic segment. **F-J.** Analysis of excitatory synapses onto engram cells. **F.** Representative images of BA of Linked and Not Linked group mice showing engram B dendrites (RFP signal, pink), VGLUT1 (red) and PSD- 95 (green). DAPI signal is represented in blue. Scale bars: 10 µm, insets: 7.5 µm. **G-J**. Analysis of excitatory synaptic markers in Linked and Not Linked animals 45 min after recall of A’. **G.** Total number of excitatory synaptic contacts onto engram cells measured as number of VGLUT1-PSD-95 doublets onto engram B - see materials and methods for more details. **H.** Number of VGLUT1^+^ puncta that are located less than 1 µm distant to an engram surface. **I.** Number of PSD-95^+^ puncta in engram cells is defined as the total number of PSD-95^+^ puncta that are located less than 1 µm distant from the surface of engram cells. **J.** Total PSD-95 expression in engram cells is quantified as the average mean intensity of PSD-95 signal inside engram cells. **K-O.** Analysis of inhibitory synapses onto engram cells. **K.** Representative images of BA of Linked and Not Linked group mice showing engram B dendrites (RFP signal, pink), VGAT (red) and GPHN (green). DAPI signal is represented in blue. Scale bars: 10 µm, insets: 5 µm. **L-M.** Analysis of inhibitory synaptic markers in Linked and Not Linked animals 45 min after recall of A’. **L.** Total number of inhibitory synaptic contacts onto engram cells measured as number of VGAT-GPHN doublets onto engram B – see materials and methods for more details. **M.** Number of VGAT^+^ puncta that are located less than 1 µm distant to an engram surface. **N.** Number of GPHN^+^ puncta in engram cells is defined as the total number of GPHN^+^ puncta that are located less than 1 µm distant from the surface of engram cells. **O.** Total GPHN expression in engram cells is quantified as the average mean intensity of GPHN signal inside engram cells. Data presented as mean ± SEM. n = 7 - 8. Nested ANOVA, Sidak multiple comparisons (C-E); unpaired t-test; *p<0.05; **p<0.01; ns, not significant. GPHN, Gephyrin.

Age has been associated with reduced synaptic plasticity mechanisms ^43,44^. We investigated whether aged mice were able to update the memory, and 9 - 11 month old mice showed no deficits in recall of a memory of an updated context, nor difficulties in recognizing the neutral or fearful contexts (Suppl Fig 6).

### Engram specific manipulation of PSD-95 modifies synaptic wiring and impairs extinction of a memory

In order to investigate the role of PSD-95 in engram synaptic wiring, we knocked-down the expression of PSD-95 specifically in the postsynaptic BA component of the B engram with a cocktail of three shRNAs. We locally injected the cocktail AAV_9_-c-fos-tTA and AAV_9_-TRE- shRNADlg4-RFP vectors into the BA (Fig 6, A). After validating the manipulation by qPCR in the BA (Suppl Fig 7), both PSD-95 knock-down and control (Scramble) animals went through the memory updating behavioural protocol (Fig 6, B). While animals were able to update A’ (Fig 6, D), the knock-down of PSD-95 in engram B induced a higher level of reactivation of engram B after recall of A’ (Fig 6, E - G).

**Figure 6.**
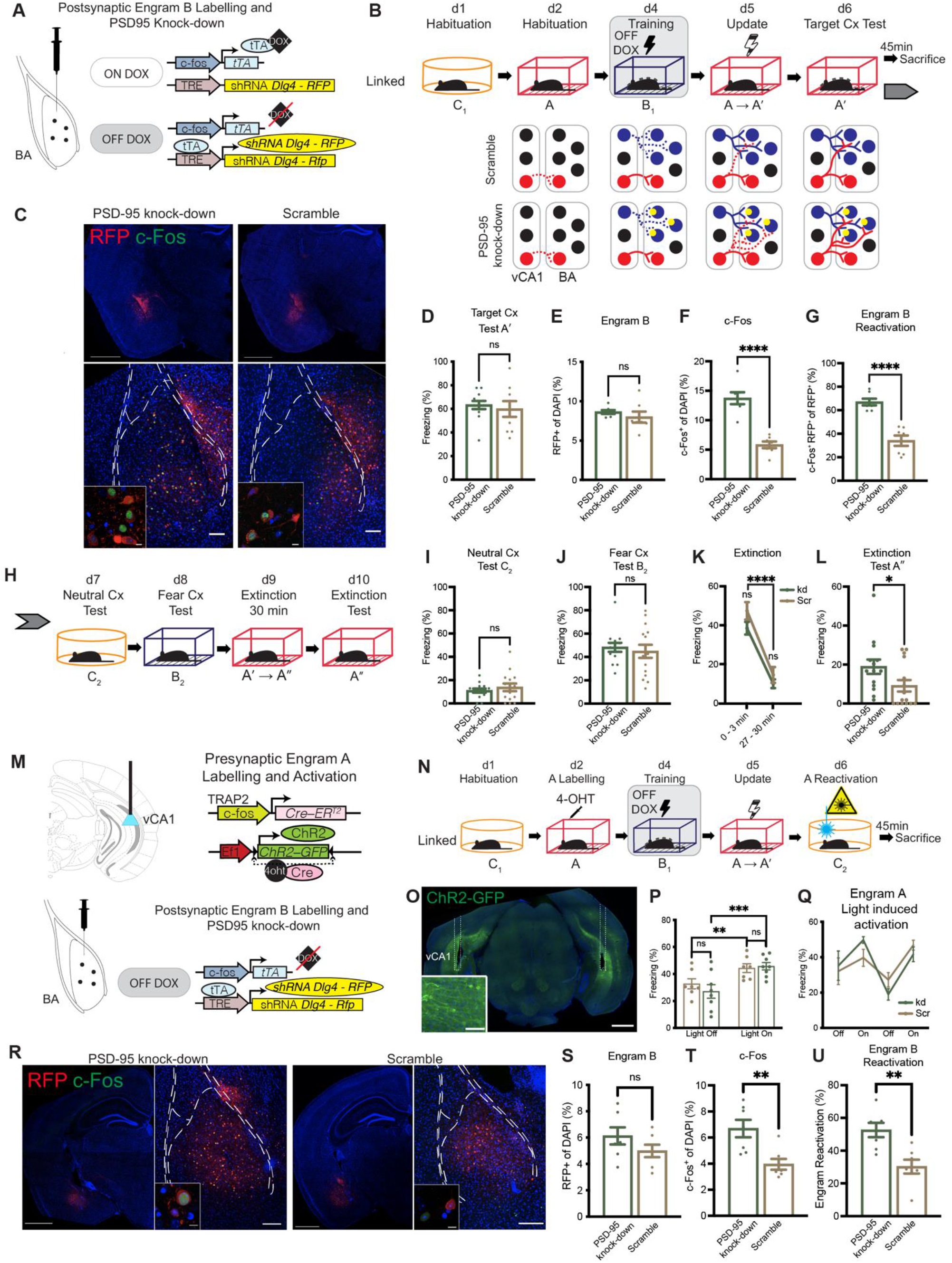
Engram specific manipulation of PSD-95 modifies synaptic wiring and impairs extinction of a memory. A. Doxycycline-dependent engram labelling strategy to knock-down Postsynaptic density protein 95 (PSD-95) while labelling engram B with a red fluorescent protein (RFP) in the BA. **B.** Behavioural schedule. After habituation to context C and A on days 1 and 2 respectively, doxycycline was removed from the diet and engram B was targeted during training on day 4 (contextual fear conditioning paradigm, black lightning symbol), with RFP and a cocktail of shRNAs anti-*Dlg4*, gene that codifies for PSD-95 (PSD-95 knock-down group, kd, yellow circles in the diagrams) or with a Scramble control (Scramble, Scr). On day 5, engram A was updated to A’ (immediate shock paradigm, black lightning symbol). On day 6, engram B reactivation induced by A’ context was measured by sacrificing mice after a natural recall of A’. **C.** Representative images of red fluorescent protein (RFP, red), c-Fos (green) and overlap of the two signals in the BA. DAPI signal is represented in blue. Scale bars: 1 mm (upper panels), 100 µm, insets: 10 µm. **D.** Knock-down of PSD-95 did not prevent updating of the memory for a neutral to fearful context A’, since PSD-95 knock-down group showed no difference in freezing compared to Scramble group. **E.** Engram B size in BA (RFP^+^ cell counts). **E.** c-Fos^+^ cell counts in BA. **F.** Higher engram B reactivation induced by natural recall of A’ in PSD-95 knock- down group compared to Scramble, indicating an aberrant higher number of connections between A and B in the absence of PSD-95. RFP^+^ and c-Fos^+^ cell counts, measured as the ratio of double positive cells per total to RFP^+^ cells. **H.** Behavioural schedule on days 7 to 10 as a continuation of behavioural schedule on panel B with a separate cohort of mice. Presence of a fear memory was tested in neutral context C (**I**) and fearful context B (**J**), showing that postsynaptic knock-down of PSD-95 did not influence recall of a neutral context C or fearful context B. **K-L.** Persistence of the updated A’ memory. Freezing decreased similarly in both PSD-95 knock-down and Scramble groups after a 30 min exposure to A’ not accompanied by any foot-shock delivery (A’ becomes A’’) (**K**). After the extinction, PSD-95 knock-down mice (kd) showed a higher retention of the updated memory than Scramble mice (Scr), which was evidenced by a higher level of freezing to A’’ (**L**). **M-U.** Optogenetic stimulation of vCA1 neutral engram cells elicits updated memory recall and higher downstream reactivation of PSD-95 depleted engram B cells. **M.** Upper panel: Tamoxifen-induced engram labelling strategy to allow optogenetic activation and tagging of engram A by AAV-delivery of a Cre-dependent channelrhodopsin (ChR2) protein and EYFP reporter into the vCA1 of TRAP2 mice. Optic fibers were implanted to allow the delivery of the light. Bottom panel: Doxycycline-dependent engram labelling strategy to knock-down postsynaptic density protein 95 (PSD-95) while labelling engram B with a red fluorescent protein (RFP) in the BA. **N.** Behavioural schedule. Animals were habituated to context C on day 1. On day 2, labelling of engram A was achieved by the injection of tamoxifen (4-OHT) immediately after habituation to A. Doxycycline (DOX) was removed from the diet of the animal and on day 4 engram B was targeted and labelled during training (contextual fear conditioning paradigm, black lightning symbol). On day 5, engram A was updated to A’ (immediate shock paradigm). On day 6, the presynaptic vCA1 component of engram A was artificially activated by delivery of light (blue laser beam symbol, yellow laser symbol), while mice explored neutral context C. Engram B reactivation induced by artificial activation of A’ was measured by sacrificing mice 45 min after optogenetic stimulation. **O.** Representative images of EYFP signal (green) in vCA1 and implant location. DAPI signal is represented in blue. Scale bars: 1000 µm, inset: 100µm. **P-Q.** Light-induced activation of A’ elicited freezing while in Context C in both Scramble and PSD-95 knock-down mice. **R.** Representative images of red fluorescent protein (RFP, red), c-Fos (green) and overlap of both signals in the BA. DAPI signal is represented in blue. Scale bars: 1000 µm (left panels), 200 µm (right panels), 10 µm (insets). **S.** Engram B size in BA (RFP^+^ cell counts). **T.** c-Fos^+^ cell counts in BA. **U.** Engram B reactivation induced by optogenetic recall of A’. RFP^+^ and c-Fos^+^ cell counts, measured as the ratio of double positive cells per total to RFP^+^ cells. Data presented as mean ± SEM. n = 9 - 12 (D); n = 7 (E-G); n = 14 - 15 (I - L); n = 7 – 8 (P, Q); n = 7 (S - U). Unpaired t-test (D – G, I – J, L) or Repeated Measures two-way ANOVA, Sidak multiple comparisons (K, P); *p<0.05; **p<0.01; ****p<0.0001; ns, not significant. BA, basal amygdala; Cx, context; DOX, doxycycline.

This functional enhancement of the connection between A and B did not modify the number of dendritic spines (Suppl Fig 8), nor the response to a neutral context (Fig 6, H, I) or the recall of fearful context B (Fig 6, J). However, we explored whether this re-wiring had effects on the persistence of the updated engram. 30 min exposure to A’ decreased freezing in both groups (Fig 6, K), whereas PSD-95 knock down mice retained the fear memory to A’ when compared to Scramble mice (Fig 6, L).

To further analyse the effect of depletion of PSD-95 in synaptic wiring, we combined the injection of the shRNA cocktail in the postsynaptic component in the BA with a presynaptic optogenetic tagging strategy. We labelled the presynaptic engram cells of the connection (vCA1) during the neutral-context encoding with a locally injected AAV_9_-DIO-ChR2-GFP vector into TRAP2 animals ^38,39^ (Fig 6, M - O). After updating A context to A’, artificial activation of A’ by optogenetics in a neutral context C was sufficient to elicit a fear response (Fig 6, P – Q) and mimic the levels of engram B reactivation in the knock-down induced by natural recall of A’ (Fig 6, R – U). Taken altogether, these results suggest that the lack of PSD-95 results in an over-reactivation of engram B that could be explained by an excess of functional connectivity between these two ensembles.

## DISCUSSION

Although much is known about how engram cells contribute to memory function at a behavioural level, less is known about the cellular and molecular mechanisms that enable engram cell plasticity. We demonstrate here that experiences are translated into changes in engram connectivity patterns across a monosynaptic connection, and that these changes are required for the recall of the updated memory. Finally, we identify a requirement for PSD-95 in engram-specific synaptic wiring.

This study sheds light onto the problem of how information specificity is achieved in the brain. We demonstrate here that a single experience reroutes the engram to recruit new ensembles and by doing so, it forms a meaningful memory. Our results sustain the hypothesis that information can be encoded in the connectivity pattern between engram cells, that adapts in parallel to animal behaviour ^9^, survives some forms of amnesia ^10,26–31^, and can be maintained long-term through cellular and synaptic homeostasis ^10,15,17,18,24,25^. Other studies have pointed in the same direction, indicating that the activation of downstream ensembles is required for memory recall ^9,10,45–47^, characterising how brain-wide distributed engrams are functionally able to elicit recall of a particular memory ^23,48–52^ or generating artificial memories and engram associations that involve interconnectivity between different areas ^8,32^. However, by studying a defined monosynaptic connection that serves a single component of the circuit, we are able to monitor the synaptic change in wiring through a behavioural readout, demonstrate that it changes to mirror experience, pinpoint a mechanism that participates in that change, and offer a paradigm to further analyse those changes.

Different areas within the BA have been associated with positive and negative valence encoding ^53^. Locally segregated in the BA, they exhibit different afferent connections ^54^. It is therefore reasonable that, as we describe here, the engram needs to be rewired from neutral to aversive populations in the BA. On the other hand, encoding of valence-associated information has also been also described in the vCA1, in two segregated subsets of cells that correlate with positive and negative experiences ^55^. In our study, it is conceivable that during the updating of the context, some negative-associated neurons in the vCA1 become at least partially recruited to the ensemble, while the previously-neutral now fear–encoding ensemble, modifies its wiring pattern.

The strengthening of a subset of vCA1-BA synapses has been suggested to contribute to adaptive fear memory in a contextual fear conditioning paradigm ^22^. We show here that by forcing the previously-neutral vCA1 engram to be updated, the synapses vCA1-BA target a different postsynaptic component, and adapt its synaptic wiring. The change in synaptic wiring we observe in our preparation could be partially derived from the structural plasticity changes reported. Similarly, projections from the vCA1 to other areas, such as the lateral hypothalamic area (LHA), have been associated with anxiety and avoidance ^56^. It is plausible that, on top of the vCA1-BA rewiring we observe, the engram connectivity pattern between vCA1 and other areas, such as the LHA, also changes after learning.

It is worth noting that the high level of dendritic spines observed in engram cells after linking does not correlate directly with histologically defined PSD-95 or SAP102 containing synapses. Our analysis does not have the technical resolution sufficient to distinguish synapsis formed by engram-engram cell connections from the broader range of synapses that these neurons receive, including from non-engram cells, or other engram cells not reactivated by that experience. That level of resolution could potentially be achieved by techniques such as dual-eGRASP ^19,20,57^, that combine activity-dependent systems with synapse tagging to label synapses between engram cells. Secondly, the membrane-associated guanylate kinase (MAGUK) family of proteins that are located at the postsynaptic area are functionally redundant and can be modulated by several interacting partners ^40,58^. It is possible that assessing the contribution of other MAGUK such as PSD-93, SAP97 or SH3 and multiple ankyrin repeat domains proteins (SHANKs)^59,60^ could help clarify this issue. Despite these caveats and limitations, our analysis show that PSD-95 expression, as well as inhibitory connections, are modulated in engram cells after experience. Therefore, studying the activity-dependent changes of postsynaptic markers is a viable method for investigating engram plasticity mechanisms.

Inhibition of engram cells has been proposed as a mechanism to explain perception and memory ^61^. Our results show downregulation of the GABA receptor clustering protein gephyrin in linked engram cells. This result is in agreement with other studies, that found a transcriptional downregulation of gephyrin in the BA 2h after contextual fear conditioning ^62^, as well as a decrease in the amount of GABA released in the BA after recall of a fear memory in an auditory fear memory paradigm ^63^. A reduction of this inhibitory marker on engram cells after they have been updated with fear information could contribute to the mechanistic explanation of why such an engram is more prone to be recalled upon appropriate fear-associated cues.

Extinction is considered a particular type of adaptive learning, in which new and relevant information, such as the lack of aversive stimulus presentation in a fearful context for a significant duration of time, *suppresses* or *modifies* the previous information and influences future behaviour ^3,64^. Mechanistically, both the *suppression*, by recruitment of new extinction-engrams ^65–67^, as well as *modification*, by weakening of the original synapses ^68^ have been demonstrated to be associated with this process. The extinction deficit shown by the decrease in PSD-95 in this study can be both interpreted as a *suppression* problem caused by an imbalance between the original fearful engram neurons that outnumber the extinction engram or as a *modification* consequence of a higher number of connections that could be therefore more resistant to weakening. Alternatively, a third explanation could be that the lack of PSD-95 of the original engram influences the interaction with other areas implicated in extinction, such as the medial prefrontal cortex ^69^.

PSD-95, by interacting with NMDA and AMPA receptors, regulates synaptic plasticity in response to activity, and training in a context preference paradigm downregulates PSD-95 expression ^70^, which resonates with what we found in our study of engram cells. PSD-95^+^ puncta decrease their size after training animals in an spatial-choice task, as well as the total expression of the protein decreased ^70^. Regulation of PSD-95 has also been shown to be required to extinguish a fear memory. For example, animals constitutively lacking a functional PSD-95 protein were able to encode and recall recent fear memories in a tone-cued paradigm, but were unable to extinguish them ^71^. Repeated exposure to the tone in the absence of a foot-shock did not decrease the extinction-resistant fear memory formed in the absence of functional PSD-95. Interestingly, this knock-out model showed impaired context discrimination where animals exhibited fear generalization to other neutral contexts. In our PSD-95 engram-specific depletion model, however, fear generalization does not occur, which is evidenced by the fact that the effects of the lack of PSD-95 do not expand to other neutral contexts. Similarly, PSD-95 has been demonstrated to be regulated by learning. Extinction of a contextual fear conditioning memory modulated expression of PSD-95 in dorsal CA1 (dCA1), increasing the protein levels at the dendritic spines and decreasing the overall expression levels ^72^. Furthermore, extinction also manifested in a phosphorylation of the PSD-95 protein, and local genetic ablation of such phosphorylation in dCA1 was sufficient to prevent memory extinction ^72^. Finally, depletion of NMDAR function induced by the local genetic deletion of the *Grin1* gene by cell transfection of a AAV-CRE into a *loxP* flanked model in dCA1 did not impair learning but induced fear extinction attenuation ^73^. NMDAR-deficient animals failed to extinguish the association between tone and shock in a cued-contextual fear paradigm. Similar to our study, mutant NMDAR animals showed a reduction in their freezing response during the extinction session, followed by a higher level of freezing to the tone on the next day. Similarly, in a contextual conditioning paradigm, repeated exposure to the context was not sufficient to extinguish the fear memory in this local NMDAR depletion model. Our study suggests that these synaptic changes mediated by PSD-95 are likely happening in engram cells, and therefore showing an effect on engram synaptic wiring.

In our model, synaptic rewiring seems to be favoured by the reduction of PSD-95. Mutant mouse models that result in functional PSD-95 ablation such as the PSD-95 knock-out model ^35^, or the brain-specific angiogenesis inhibitor 1 (BAI1) knock-out mice ^74^ result in an increase in long-term potentiation in the hippocampus, while long-term depression is absent ^35^. LTP enhancement has been associated with a preferential allocation of memories to ensembles ^75^, and therefore it is reasonable to speculate that the PSD-95-deficient engram B will be preferentially rewired during memory updating in our paradigm.

In conclusion, we have described how plasticity of synaptic wiring is a credible candidate mechanism that allows engram cells to be updated with new experiences, influencing memory function. We also describe a cellular mechanism based on PSD-95 that acts as a bridge to translate synaptic plasticity changes into synaptic wiring changes. Future research is needed to clarify which are the synaptic changes that, influenced by PSD-95 depletion, control the wiring of engram cells. Overall, this study helps us understand the role of engram cells in learning and memory, by offering a feasible conceptual explanation for memory storage and memory function.

## ACKNOWLEDGEMENTS

We thank Lydia Marks and Mariia Yurova for technical and administrative support and Elizabeth Dura for experimental assistance. We thank Tamara Boto and all the past and present members of the Ryan Lab for support and discussion. We thank Livia Autore, Andrea Muñoz Zamora, Erika Stewart and Aaron Douglas for proofreading. This work was funded by the Irish Research Council (GOIPD/2019-812), European Research Council (715968), Science Foundation Ireland (15/YI/3187) and the National Institute of Health (1R01NS121316).

## AUTHOR CONTRIBUTIONS

C.O.S. and T.J.R. conceived the scientific design. C.O.S. conducted the experiments and analysed the data. E.U. performed experimental work. M.P. conducted electrophysiological characterizations. C.O.S. and T.J.R. interpreted the results. C.O.S. and T.J.R. wrote the original draft. C.O.S., M.P., and T.J.R. reviewed and edited the final manuscript.

## MATERIALS AND METHODS

### Animals

The experimental subjects employed in this study were male C57Bl6/J mice (Charles River), aged 7 to 14 weeks, or between 40 and 48 weeks for the ageing experiments. TRAP2 mice (Fos^tm2.1(iCre/ERT2)Luo^/J, JAX stock #030323) ^38,39^ were used to achieve double engram labelling. The animal room was maintained at a consistent temperature of 22°C, and a 12-hour light/dark cycle was established, with all experimental steps conducted during the light phase. Mice were housed in groups of five in cages equipped with a tunnel, with free access to food and water. The care and behavioural experiments involving the mice adhered to the guidelines provided by the Animal Ethics Committee of Trinity College Dublin, the Health Products Regulatory Authority of Ireland, and the European Directive 2010/63/EU.

### Engram labelling

Under anaesthesia induced by administering 500 mg/kg of avertin, the mice were positioned with their heads fixed onto the stereotactic frame (WPI). Bilateral craniotomy was performed on each animal with an electric drill at appropriate injection coordinates (0.5 mm diameter drill, WPI). The viral mix was delivered using a 10 μL Hamilton syringe (701LT; Hamilton) and a microsyringe pump (UMP3; WPI). The injection speed was 60 nL/min. Cocktail AAV volumes injected and injection coordinates with respect to Bregma were as follows: BA, 250 nL of AAV, injected at 20 degrees angled, anteroposterior (AP) +0.45 mm, mediolateral (ML) ±3.00mm, dorsoventral (DV) -5.5mm; vCA1, 350 nL, AP -3.5, ML ±3.5, DV -3.8. AAVs used were: Fig 2, 3, 5, Suppl Fig 2, Suppl Fig 3, AAV_9_-c-fos-tTa (1:10, gift from S. Tonegawa, AA_9_-TRE-RFP (6:10, gift from S. Tonegawa); Fig 3, 6, AAV_9_-EF1a-DIO-hChR2-EYFP (1:2, Addgene viral prep # 20298- AAV9; http://n2t.net.addgene:20298; RRID; Addgene:20298, gift from Karl Deisseroth); Fig 4, AAV_9_-c-fos-tTa (1:10), AAV_9_-TRE-ArchT-EYFP (4:10, gift from S. Tonegawa); Fig 6, Suppl Fig 5 AAV_9_-c-fos-tTa (1:10), AAV_9_-[miR30]-TRE>RFP:{mDlg4[miR30-shRNA#1,2,3/or Scramble]}:WPRE] (1:3 for shRNA#1,2,3 in a cocktail; 1:3 Scramble; Custom design by VectorBuilder); Suppl Fig 3: AAV_9_-CaMKIIa-hChR2-EYFP (1:5, Addgene viral prep #26969- AAV9; http://n2t.net.addgene:26969; RRID; Addgene:26969, gift from Karl Deisseroth; AAV_9_- c-fos-tTa (1:2), AAV_9_-TRE-ChR2-EYFP (1:5, gift from S. Tonegawa). All the AAVs, except when otherwise stated, were packed by Charles River Laboratories into a titre of at least 10^13^GC/mL. Additionally, in optogenetic experiments, two fiber implants (200 µm core diameter, Doric lenses) were affixed to the stereotaxic adapter and carefully lowered into the brain above the viral injection site: vCA1, AP -3.5, ML ±3.5, DV -3.7; BA, AP -1.55, ML ±3, DV -4.6. Implants were securely adhered using adhesive (C&B Metabond) and dental cement (Dentalon plus, Kulzer). Mice were administered analgesia (meloxicam, 1.5mg/kg, subcutaneously) during the following two days, and allowed a recovery period of 10 days prior to the commencement of any behavioural procedures.

Engram labelling is based on the expression of transgenes controlled by the promoter of immediate early genes, such as *c-Fos* ^5^ temporally regulated by the presence of doxycycline (DOX) or tamoxifen ^5,39^. To label the engram in DOX dependent strategies, mice were fed DOX-enriched food (40mg/kg, Datesand) from at least 24 h prior to surgery and maintained on it for the duration of the protocol except for the labelling window. DOX was removed from the diet for 36 hours prior to the target experience to label. In the case of labelling a random ensemble (Figure 4), DOX was removed from the diet for 50 hours in total and during that period, homecage was changed twice, new nesting material and homecage enrichment (tubing) was introduced, as well as manipulation by the experimenter. Immediately after experience, mice were fed again with the DOX diet. To label the engram in tamoxifen dependent strategies, 4-hydroxytamoxifen (4-OHT, Santa Cruz) was freshly prepared in oil (mix of sunflower seed oil, castor oil, Sigma; 4:1) at 10 mg/mL ^38^ and administered through intraperitoneal injection (50 mg/kg) right after the experience.

### Behaviour

Three different contexts were used. Context A was a 31 x 24 x 21 cm arena (Med Associates) equipped with a removable grid floor (3.2mm diameter), opaque triangular inset ceiling, scented with benzaldehyde (0.25% in Ethanol 70%, Sigma), and illuminated with maximum white light intensity. Context B was a 29 x 25 x 22 cm arena (Coulbourne Instruments), with a grid floor, scented with acetic acid (1% in H_2_O, Sigma) and medium light levels. Context C was a 40 x 20 x 30 cm arena with a grey floor and grey walls. Each of the contexts was located in a different behavioural room, and mice were brought into the room in a transportation box. To habituate mice to the experimenter and the transportation box, mice were handled for 5 minutes per animal and habituated to the transportation box once a day during 3 days prior to any behavioural protocol. Timelines for the behavioural protocols are specified in the schematics included in figures. During habituation to the contexts C and A, mice were allowed to freely explore the contexts for 10 minutes. During training, mice underwent a contextual fear conditioning training paradigm, in which they were allowed to explore the arena for 3 minutes, after which they were presented with 3 foot shocks (0.2 s, 0.75mA) spaced by intervals of 1 min. For the updating, mice were introduced in the arena and immediately received a single foot shock (2 s, 1.25mA). They were removed from the arena 1 min after. In the control group that did not receive an update, mice were allowed into the arena for 1 min but no foot shock was delivered. During the tests, mice were exposed to the contexts for 3 min and their freezing behaviour was scored. During the extinction protocol, mice were allowed to explore the arena for 30 min.

### Optogenetics

The optogenetic activation of vCA1 was achieved by light delivery into vCA1, with a 450 nm diode fiber light source (Doric Lenses). Light was bilaterally delivered (4 Hz, 6-9mV) during 3 minute episodes initiated at time 3 min and time 9 min. Session was 12 min long. The optogenetic inhibition of BA was achieved by light delivery into BA, with a 520 nm diode fiber light source (Doric Lenses). Light was bilaterally delivered (20 Hz, 6-9mV) during the 3 min long session in Context A.

### Histology

Mice were intraperitoneally anaesthetised (Sodium Pentobarbital, Eutanimal) and transcardially perfused with phosphate buffered saline (PBS, Sigma) and paraformaldehyde (PFA-4% in PBS, Sigma). Brains were extracted and were postfixed by immersion in PFA-4% in PBS at RT overnight. 50 µm coronal brain slices were obtained with a Vibratome (Leica) and immunostained. For c-Fos overlap analysis, mice were perfused 45 min after behaviour was finalised (natural recall or optogenetic reactivation). To perform overlap analysis, tissue was permeabilized in PBS-Triton 0.2%, blocked in Normal Goat Serum 10% - Triton 0.2% in PBS for 1 h at RT, shaking, and incubated with anti–c-Fos antibody (1:1000, Synaptic Systems 226- 008) in blocking solution overnight at 4 degrees. After washes, tissue was incubated with anti-rabbit Alexa-488 (1:800, Invitrogen, A11034) for 2 hours at RT. Tissue was further washed, incubated in DAPI (1:1000, Sigma, D9564) for 5, washed in PBS and mounted (Vectashield, Vector Laboratories). For synaptic markers analysis and dendritic spine density analysis, tissue was immunostained following the same protocol with primaries anti-PSD-95 (1:500, Invitrogen, MA1-045), anti-VGLUT1 (1:400, Synaptic Systems, 135-304), anti-VGAT (1:1000, Synaptic Systems, 131 005), anti-GPHN (1:500, Synaptic Systems, 147 008) and anti-RFP (1:1000, Rockland, 600-401-379). In the case of anti-SAP102 staining (1:250, Neuromab, 75- 0558), tissue was subjected to an antigen retrieval protocol consisting in an incubation with boiling sodium citrate buffer (10 mM, 0.05% Tween-20, pH 6.0) during 30 min prior to permeabilization. Secondaries used were anti-mouse Alexa 488 (1:800, Invitrogen, A11001), anti-rabbit Alexa 568 (1:800, Invitrogen, A11011) and anti-guinea pig AlexaFluor AffiniPure 647 (1:800, Jackson ImmunoResearch, 106-605-003). To detect ArchT-EYFP and ChR2-EYFP, tissue was immunostained with primary anti-EGFP (1:1000, Invitrogen, A10262) and secondary anti-chicken Alexa 488 (1:800, Invitrogen, A11039).

### Imaging and images analysis

Images were obtained using a Leica SP8 confocal microscope at 40X magnification. Confocal microscope settings for imaging were maintained constant between individuals. For cell counting, single plane images were obtained from 3-6 sections and values were averaged per animal. Images were analysed using ImageJ (NIH) and Photoshop (Adobe). Total number of engram positive and c-Fos^+^ cells were manually counted per area, whereas total number of DAPI positive cells was estimated based on area and cell density on the area. For marker analysis and spine morphology analysis, 8 µm stacks (HC PL APO 40X/1.30 OIL, Pinhole 1.00AU) were analysed using IMARIS software (Imaris v9.5, Oxford Instruments). The dendrites of BA engram cells were semi-automatically traced and spines were identified. Spine volume, spine head diameter and spine density parameters were measured by the software in 10x10 µm independent dendritic segments per animal. Dendritic spine morphology analysis was done with a semi-automated classification method, based on previous studies^76^. Stubby spines were defined as spines with less or equal to 1 µm length, and the ratio between the spine volume head and the spine volume neck was minor to 1.2. Mushroom spines were defined as spines with less or equal to 5 µm length and the ratio between the spine volume head and the spine volume neck was higher or equal to 1.2. Thin spines were defined as spines with less or equal to 5 µm length and the ratio between the spine volume head and the spine volume neck was less than 1.2. Spines where no neck was identified were classified as stubby. Synaptic puncta analysis was performed using IMARIS automatic spot detection and averaged per volume of engram cells. Number of synaptic contacts onto engram cells was calculated by first identifying the number of PSD-95^+^, GPHN^+^ or SAP102^+^ puncta inside the volume drawn over engram cells that were also in between -1 µm and 0 µm distant to the surface of engram cells. This value was represented as PSD-95^+^, GPHN^+^ or SAP102^+^ puncta in engram cells. Then, VGLUT1^+^ or VGAT^+^ puncta outside engram cells were calculated by first identifying the number of VGLUT1^+^ or VGAT^+^ puncta outside the volume drawn over engram cells, and then, selecting those that were in between 1 µm and 0 µm distant to the surface of engram cells. This value was represented as VGLUT1^+^ or VGAT^+^ puncta into engram cells. The ratio of synaptic contacts into engram/non engram cells was calculated as the percentage of VGLUT1^+^ contacts onto engram cells respective of total contacts in the region of interest (ROI) imaged. Finally, the number of synaptic contacts onto engram cells was calculated as number of doublets formed by the presence of a VGLUT1^+^ puncta into engram cells (0-1 µm apart from the surface) and a PSD-95^+^ puncta in engram cells (0-1 µm apart from the surface) separated by a distance of 0 µm to 1.8 µm (synaptic contacts onto engram cells), by the presence of a VGLUT1^+^ puncta/SAP102^+^ puncta separated by a distance of 1.8 µm, or by the presence of a VGAT^+^ puncta/GPHN^+^ puncta separated by a distance of 0 µm to 1.12 µm (inhibitory synaptic contacts onto engram cells). These synaptic markers parameters were then normalised per total volume of engram cells measured in the ROI, and 2 to 4 ROIs were averaged per animal to obtain each datapoint. Finally, the total amount of PSD-95, SAP102 or GPHN expression inside engram cells was calculated by generating a mask with the surface drawn over engram cells using IMARIS, and obtaining the average mean signal intensity within the volume inside engram cells.

### qPCR

Animals were sacrificed by cervical dislocation. Brain was quickly extracted in RNAse free conditions, immersed in RNAse free cold PBS (Invitrogen), and amygdalar area was dissected under a scope and immediately flash frozen in liquid nitrogen. Tissue was homogenized with a RNase-free glass 0.5mL Dounce and RNA was extracted with Trizol following manufacturer instructions (Invitrogen). 500ng of RNA were retrotranscribed (Quantiscript, Quiagen) following instructions from the manufacturer and qPCR was performed (iTaq Universal, Biorad). *Dlg4* mouse gene was amplified using primers Forward 5’- GCCTACCAAAGACCGTGCCA-3’ and Reverse 5’-ACTGGCCAGCCTCAATGAACT-3’. *Gapdh* was amplified as a housekeeping with primers Forward 5’-TGTGTCCGTCGTGGATCTGA-3’ and Reverse 5’-TGTCATCATACTTGGCAGGTTTCT-3’. ΔCt method was used to calculate the expression per sample as normalised per housekeeping.

### Slice preparation

Animals were anaesthetised, the brains then collected and dissected in oxygenated (95% O_2_, 5% CO_2_) cold cutting artificial cerebral-spinal fluid solution (in mM): 213 Sucrose, 2.5 KCl, 10 MgCl6H_2_O, 1.25 NaH_2_PO_4_, 0.5 CaCl_2_H_2_O, 25 Glucose, 25 NaHCO_3_. Coronal sections of 300 µm thick, containing BA were obtained with a Vibratome (Leica). Slices were collected and let to recover at room temperature for one hour, and then during the recordings maintained and controlled at 32°C (Temperature controller IIV, Luigs & Neumann) in oxygenated (95% O_2_, 5% CO_2_) artificial cerebral-spinal fluid solution (in mM): 125 NaCl, 2.5 KCl, 1 MgCl6H_2_O, 1.25 NaH_2_PO_4_, 2 CaCl_2_H_2_O, 25 Glucose, 25 NaHCO_3_. During the recordings the slices were in presence of extracellular Gabazine 10 µM.

### Ex vivo electrophysiology

Whole cell recordings in voltage clamp mode were performed under the guidance of a motorised infracontrast DGC microscope (LNScope, Luigs & Neumann) with a water immersion 40X objective, and a CCD ICX205 camera (INFINITY 2-1R). For opto-stimulation (450 nm) of the channel-hodopsin present in the neuron terminals and for detection (540- 600 nm) of mCherry fluorescent protein, a X-Cite XLED1 lamp was used for excitation, with corresponding filter cubes (Olympus). The trigger was digitally controlled as well as the light intensity of the lamp, that could range from 100% to 1%.

The recording Ag/AgCl electrodes were fabricated with borosilicate glass pipettes (P1000, Sutter Instrument) of 7 to 10 Mohms resistance and filled with the following intracellular solution (in mM): 117 Cs-Methanesulfonate, 20 HEPES, 0.4 EGTA, 2.8 NaCl, 4 Mg-ATP, 0.3 Na-GTP, 5 TEA-Cl, 10 QX314-Br, 0.1 spermine, and mannitol to adjust osmolarity to 290mOsm, pH 7.2.

The signal was amplified with a Multiclamp 700B dual amplifier (Molecular Devices), filtered at 2 kHz, digitised (20 kHz), and acquired through an ADC/DAC data acquisition interface (ITC- 18, InstruTECH) by using a custom-made software running on Igor Pro (Wavemetrics). The evoked postsynaptic currents were recorded in each neuron at -75mV.

### Data analysis and statistics

Mice were randomly assigned to each experimental group in a cage-balanced way. Animals with mistargeted implants or viral injections were removed from the analysis. Freezing behaviour was scored manually. The experimenter was blind to the experimental group and videos and images were analysed in random order. Statistical analysis was conducted on Prism 9.0 (Graphpad software). Paired or unpaired t-test, standard one or two way ANOVA followed by a Tukey post hoc test for multiple comparisons, nested ANOVA followed by a Sidak post hoc test for multiple comparisons, or repeated measures two-way ANOVA followed by a Sidak post hoc test were used as specified in figure legends. Figures were prepared with Adobe Illustrator (Adobe).

## SUPPLEMENTAL INFORMATION

**Suppl Figure 1.**
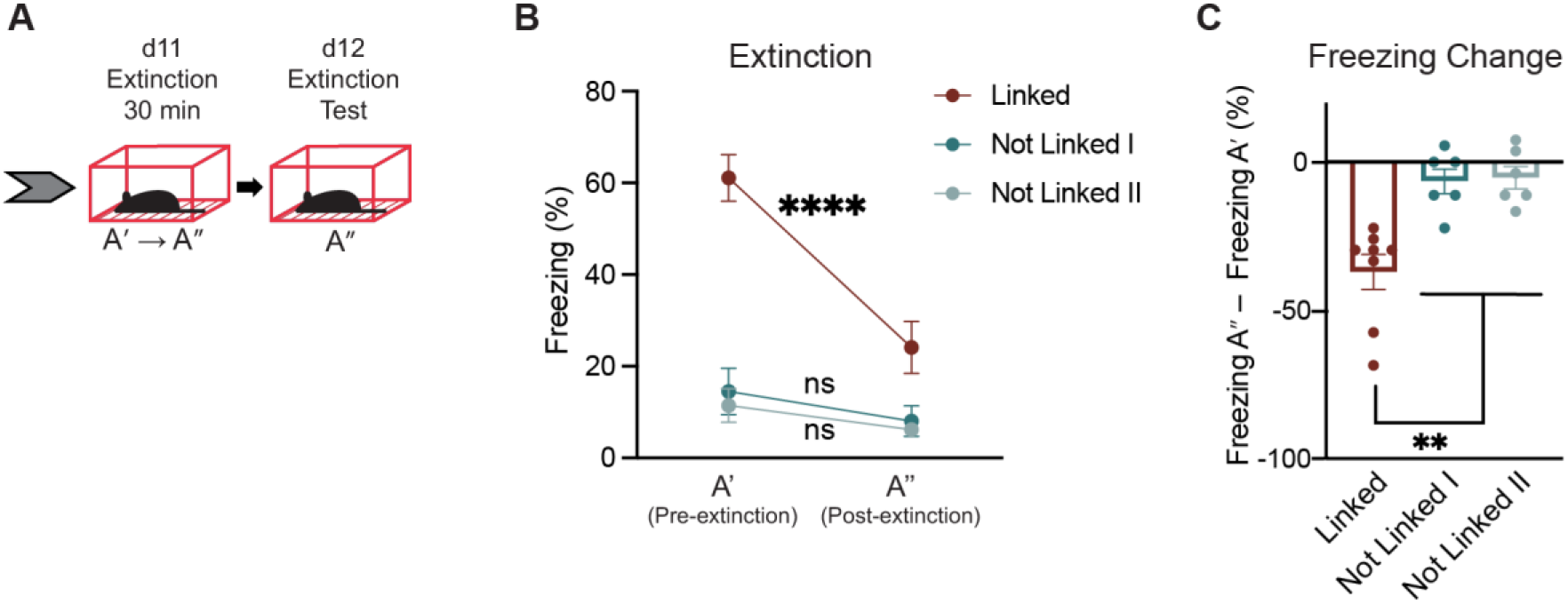
Exposure to context A’ for 30 min in the absence of footshocks induces extinction of the fear memory A. Behavioural paradigm. After experiencing the behavioural linking paradigm (Figure 1, days 1-10), on day 11, all three groups of mice (Linked, Not Linked I and Not Linked II, see Figure 1 for more details), were exposed to context A’ for 30 minutes without the delivery of any footshocks (A’ context becomes A’’). On day 12, freezing behaviour on context A’’ was tested during 3 min. B – C. Freezing behaviour is reduced in Linked animals, but not in Not Linked animals after extinction session. Freezing in A’’ test day (day 12) is significantly lower than freezing in A’ test day (day 8). C. Freezing change is calculated as the difference between freezing in context A’’ (post-extinction) and freezing in context A’ (pre-extinction). Data presented as mean ± SEM. n = 6 - 8. Repeated measures two-way ANOVA, Sidak multiple comparisons test (B), one-way ANOVA, Tukey’s multiple comparisons test (C); **p<0.01, ****p<0.0001. ns, not significant.

**Suppl Figure 2.**
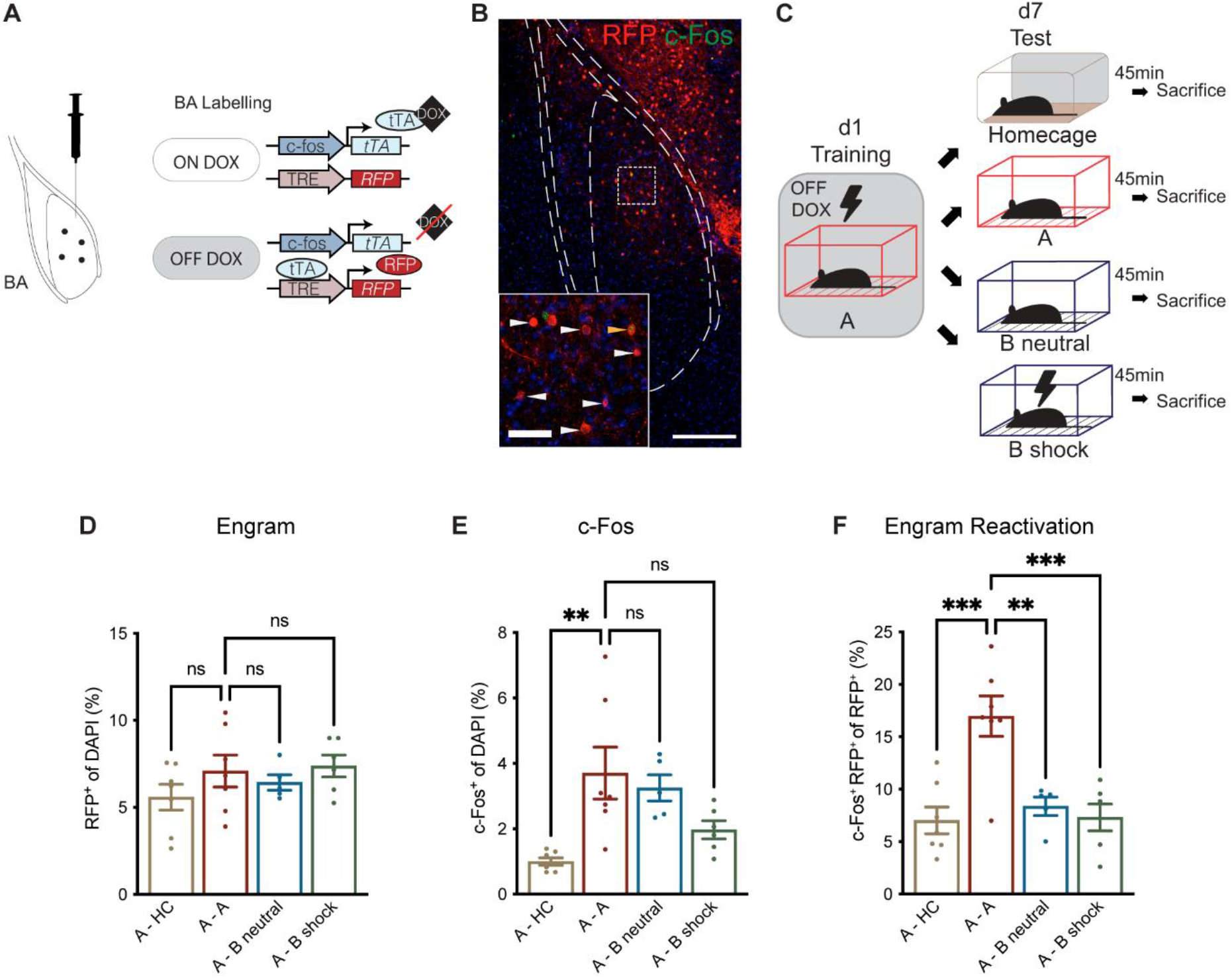
Fear memories formed by contextual fear conditioning to A and B are represented in two separate ensembles in the BA. A. Doxycycline-dependent engram labelling strategy to label engram A in the BA. B. Representative images of red fluorescent protein (RFP, red, white arrows), c-Fos (green) and overlap (orange arrow) of the two signals in the BA. DAPI signal is represented in blue. Scale bar: 250 µm, inset: 65 µm. C. Animals are trained in context A (contextual fear conditioning paradigm, black lightning symbol) on day 1, after removal of doxycycline. Fear engram for A is formed in a single experience in the BA. On day 7, engram A reactivation induced by homecage exposure, natural recall of A and B or a contextual fear conditioning training paradigm in B is measured. D. Engram A size in BA (RFP^+^ cell counts). E. c-Fos^+^ cell counts in BA. F. Engram A reactivation induced by natural recall of A is higher compared to other control groups. RFP^+^ and c-Fos^+^ cell counts, measured as the ratio of double positive cells per total to RFP^+^ cells. Data presented as mean ± SEM. n = 5 - 7. One-way ANOVA, Tukey’s multiple comparisons; **p<0.01; ***p<0.001; ns, not significant. DOX, doxycycline; HC, homecage.

**Suppl Figure 3.**
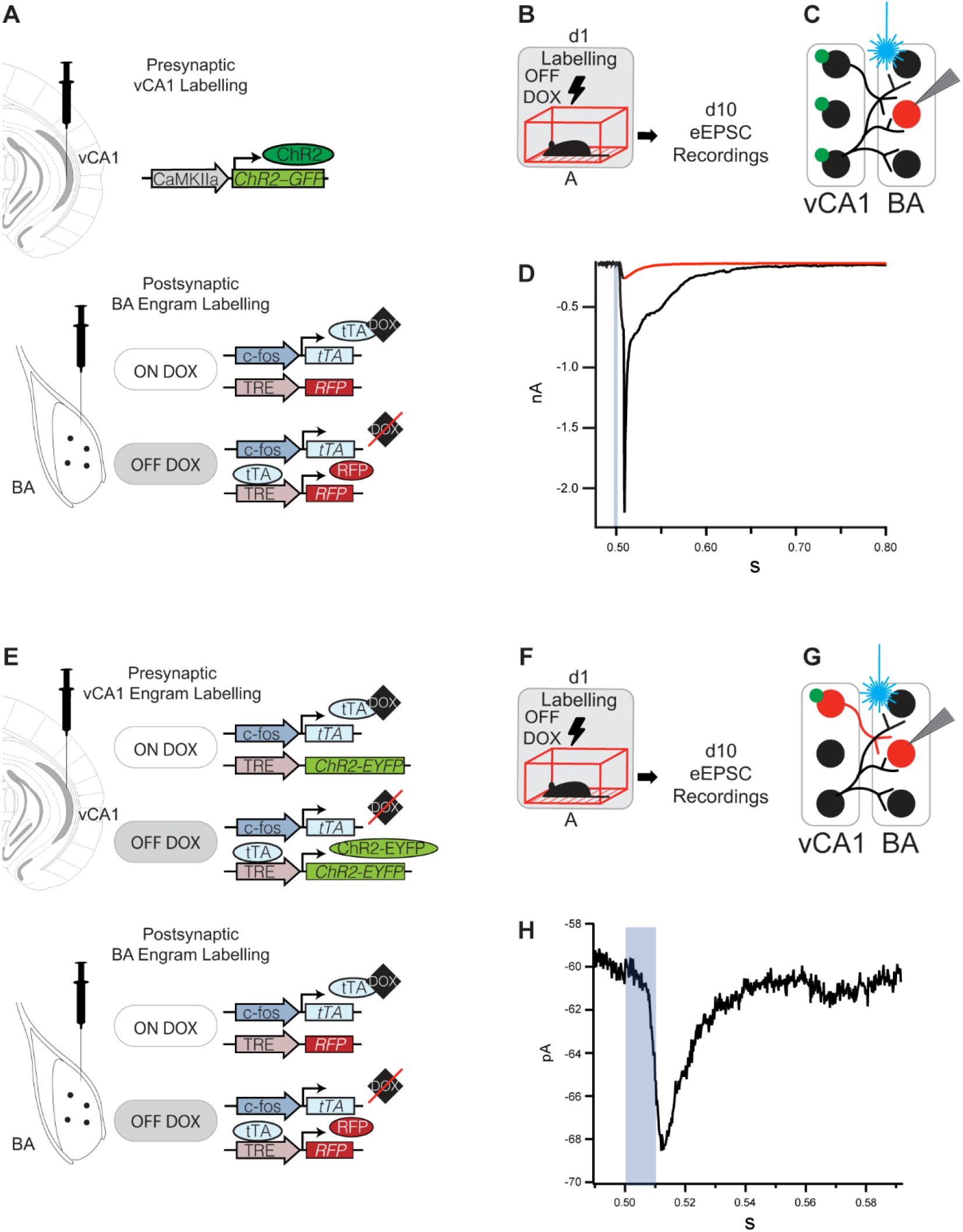
Presence of a monosynaptic connection between vCA1 and BA. A-D. Presence of a vCA-BA connection. A. Non-engram specific labelling of the presynaptic component by stereotactic injection of AAV_9_-CaMKIIa-ChR2-EYFP in vCA1 combined with the DOX-dependent engram-specific labelling of the postsynaptic component by injection of AAV_9_-c-fos-tTa and AA_9_-TRE-RFP in BA. B. Behavioural schedule. Animals were trained in Context A (contextual fear conditioning paradigm, black lightning symbol), while off DOX, on day 1. On day 10, whole cell recordings of evoked postsynaptic currents were performed. C. Schematic diagram of the connection from vCA1 to BA. Presynaptic terminals, tagged with channelrhodopsin (green circle symbol), were opto-stimulated by light-delivery (blue laser beam symbol), while response of an engram cell in the BA (red circles) was recorded. D. Average traces of evoked postsynaptic currents (eEPSCs) recorded in an engram cell in the BA after the opto-stimulation (1ms) of the ipsilateral presynaptic component terminals (blue square), with low (5%, red) and high (10%, black) light intensity. E – H. Presence of a vCA-BA connection between engram cells. E. DOX-dependent engram-specific labelling of the presynaptic component by stereotactic injection of AAV_9_-c-fos-tTA and AAV_9_-TRE-ChR2-EYFP in vCA1 combined with the DOX-dependent engram-specific labelling of the postsynaptic component by injection of AAV_9_-c-fos-tTa and AA_9_-TRE-RFP in BA. F. Behavioural schedule, similar to B. G. Schematic diagram of the connection from vCA1 to BA. Engram-specific presynaptic terminals, tagged with channelrhodopsin (green circle symbol), were opto-stimulated by light-delivery (blue laser beam symbol), while response of an engram cell in the BA (red circles) was recorded. H. Average trace of evoked eEPSCs recorded in an engram cell in the BA after the opto-stimulation (10ms, 100% light intensity) of the ipsilateral presynaptic component of engram terminals.

**Suppl Figure 4.**
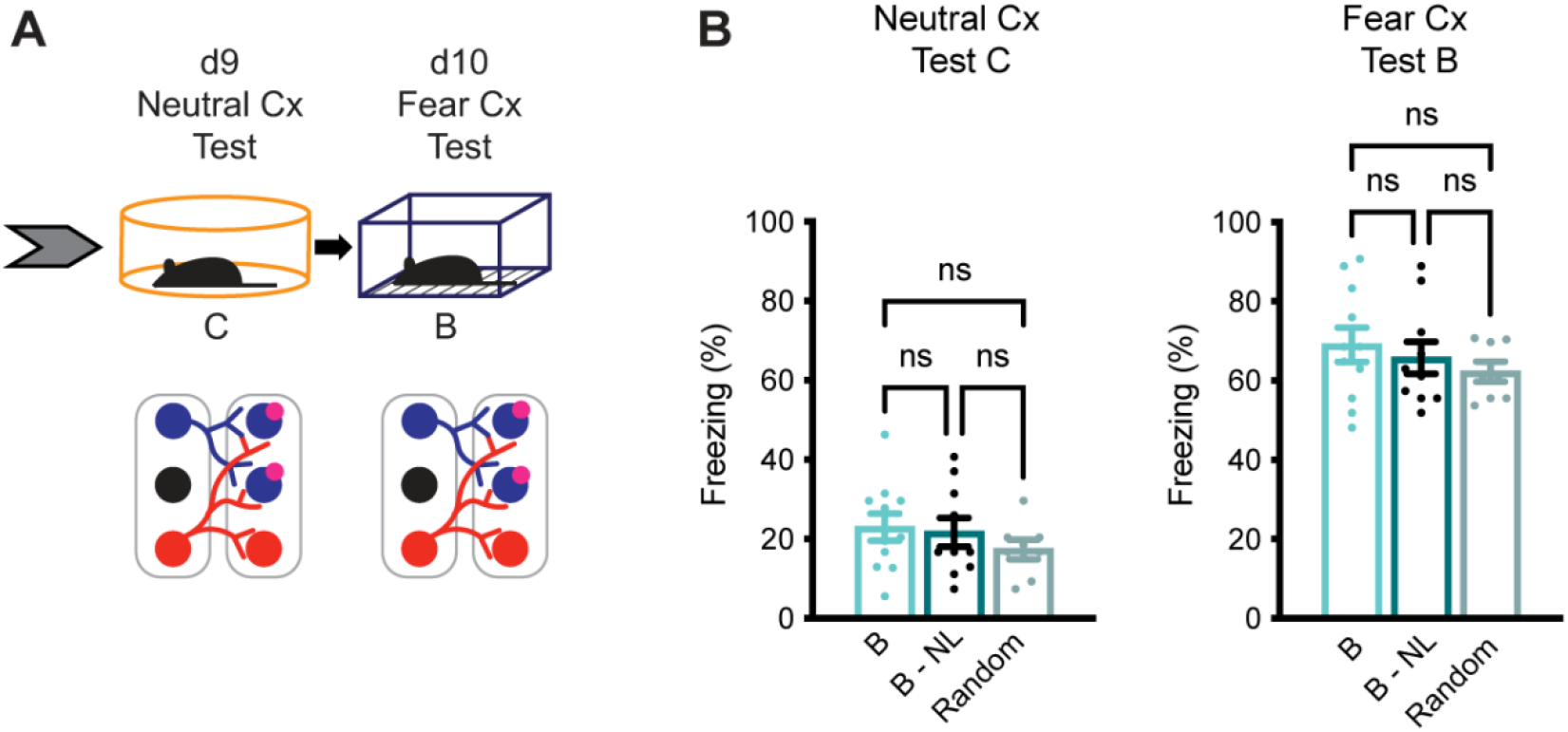
Fear response to neutral and fearful contexts not affected in mice tagged in engram B or a random ensemble. A. Behavioural schedule on days 9 and 10 as a continuation of behavioural schedule in Figure 3. Presence of a fear memory was tested in neutral context C and fearful context B. B. Freezing behaviour as a readout for memory recall of mice tagged in engram B (B and B that undergo no light stimulation during inhibition test, B – NL) is similar to those tagged in a random ensemble (Random) in a neutral context C (left) and a fearful context B (right). Data presented as mean ± SEM. n = 8 - 11. Two-way ANOVA, Tukey; ns, not significant. Cx, context.

**Suppl Figure 5.**
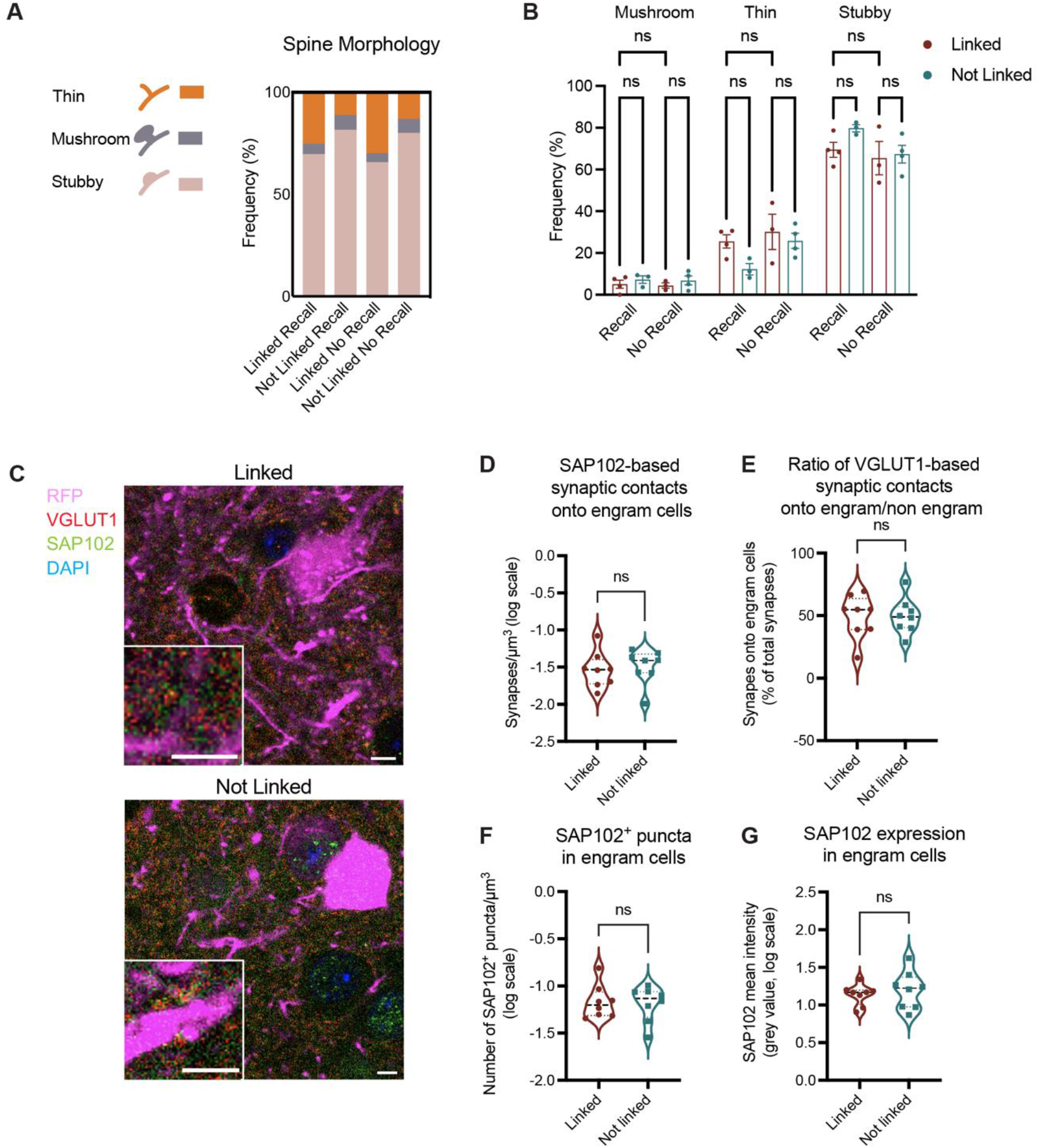
Plasticity mechanisms associated with changes in synaptic wiring in engram cells. A-B. Spine morphology characterization. Frequency of subtypes of dendritic spines in engram B cells, classified as thin (orange), mushroom (grey) or stubby (salmon). Frequency is determined by percentage of total number of spines averaged per animal in Linked (red) and Not Linked (green) groups, 45 minutes after recall induced by A’ (Recall) or homecage (No Recall). B. Comparison of the frequencies per each spine subtype. C – G. Analysis of excitatory synapses based on SAP102 postsynaptic protein in engram cells. C. Representative images of BA of Linked and Not Linked group mice showing engram B dendrites (RFP signal, pink), VGLUT1 (Red) and SAP102 (green). DAPI signal is represented in blue. Scale bars: 5 µm, 4 µm (insets). D-G. Analysis of excitatory synapses based on SAP102 in Linked and Not Linked animals 45 min after recall of A’. D. Total number of excitatory synaptic contacts onto engram cells based on SAP102, measured as number of VGLUT1-SAP102 doublets onto engram B - see materials and methods for more details. E. Ratio of synaptic contacts onto engram/non engram cells defined as the number of VGLUT1+ puncta that are located less than 1 µm distant to an engram surface, normalised by the total number of VGLUT1+ puncta. F. Number of SAP102^+^ puncta in engram cells is defined as the total number of SAP102^+^ puncta that are located less than 1 µm distant from the surface of engram cells. G. Total SAP102 expression in engram cells is quantified as the average mean intensity of SAP102 signal inside engram cells. Data presented as mean ± SEM. n = 3 - 4 animals, average of 8 - 10 x 10 µm dendritic segments per animal (A, B); n = 7 – 8 animals (D – G). Repeated measures two-way ANOVA, Sidak’s multiple comparisons test (A, B); unpaired t-test (D – G). *p<0.05; ns, not significant. BA, basal amygdala.

**Suppl Figure 6.**
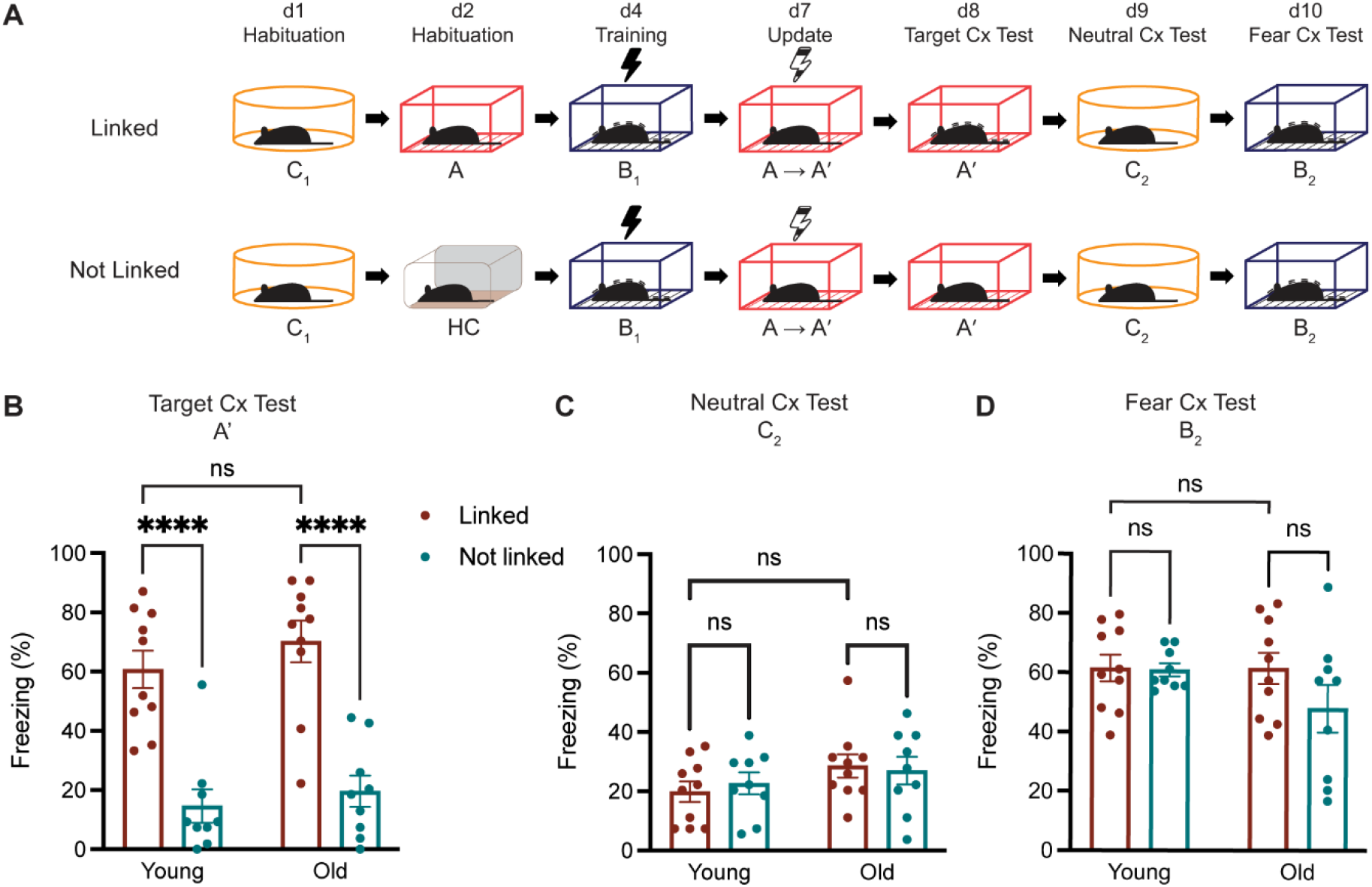
Age does not prevent updating of a neutral memory to a fearful one in old mice (9-11 months old mice). A. Aged mice (9-11 months, Old) were subjected to the behavioural schedule as explained in Figure 1 and compared to young mice (6-14 weeks, Young). B-D. Presence of a fear association to Contexts A’, C and B was tested. Old mice both in the experimental (Linked) and control group (Not Linked) showed similar freezing levels as young mice in all three contexts. Data presented as mean ± SEM. n = 9 - 10. Two-way ANOVA, Tukey; ****p<0.0001. ns, not significant. Cx, context.

**Suppl Figure 7.**
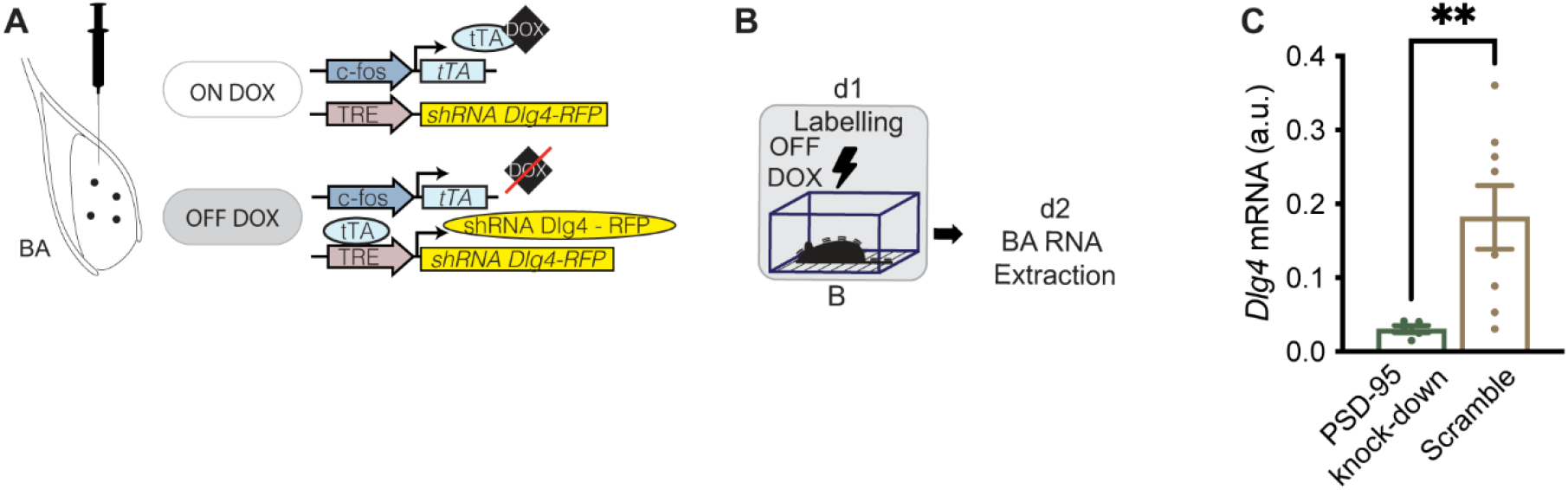
Injection of a cocktail of 3 shRNAs in a doxycycline-dependent system in the BA reduces expression of *Dlg4,* generating a PSD-95 knock-down model. A. Doxycycline-dependent system to label an engram and deplete PSD-95 in engram cells. B. Behavioural schedule. On day 1, animals were trained in context B (contextual fear conditioning, black lightning symbol) after removal of doxycycline from their diet. 24h later, animals were sacrificed and amygdalar area extracted. C. *Dlg4* (PSD-95) expression levels decreased in the PSD-95 knock-down group, injected with 3 shRNAs, when compared to the Scramble-injected group. Data presented as mean ± SEM. n = 5 - 8. Unpaired t-test; **p<0.01. ns, not significant.

**Suppl Figure 8.**
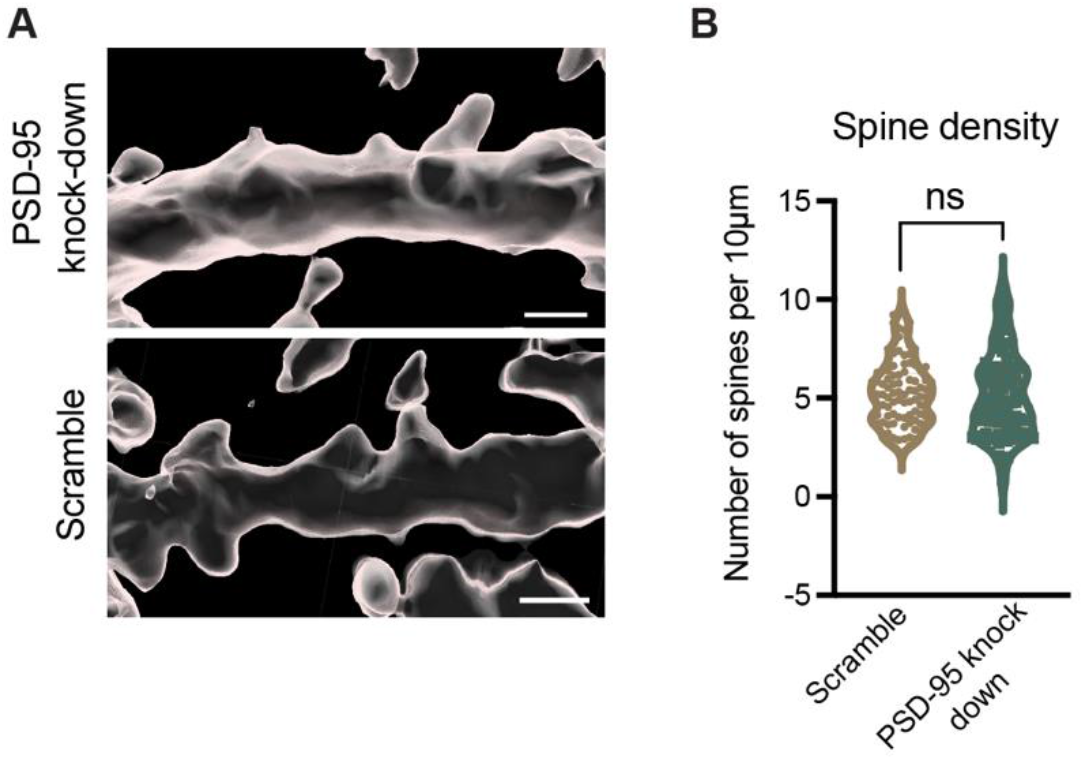
Dendritic spine analysis of engram B in animals in the absence of PSD-95. A. Representative image of a RFP^+^ dendrite as identified as a surface by the IMARIS software in each group. Scale bars: 1 µm. B. Spine density measured by number of spines in engram B cells. Data presented as mean ± SEM. n = 9 animals x 7 – 10 µm dendritic segment per animal. Nested t-test. ns, not significant.

